# Disintegration promotes proto-spacer integration by the Cas1-Cas2 complex

**DOI:** 10.1101/2020.12.21.423798

**Authors:** Chien-Hui Ma, Kamyab Javanmardi, Ilya J. Finkelstein, Makkuni Jayaram

**Author notes:** Corresponding authors: Ilya J. Finkelstein and Makkuni Jayaram, (IJF) (MJ).

## Abstract

‘Disintegration’—the reversal of transposon DNA integration at a target site—is regarded as an abortive off-pathway reaction. Here we challenge this view with a biochemical investigation of the mechanism of protospacer insertion by the *Streptococcus pyogenes* Cas1-Cas2 complex, which is mechanistically analogous to DNA transposition. In supercoiled target sites, the predominant outcome is the disintegration of one-ended insertions that fail to complete the second integration event. In linear target sites, one-ended insertions far outnumber complete proto-spacer insertions. The second insertion event is most often accompanied by disintegration of the first, mediated either by the 3’-hydroxyl exposed during integration or by water. One-ended integration intermediates may mature into complete spacer insertions via DNA repair pathways that are also involved in transposon mobility. We propose that disintegration-promoted integration is functionally important in the adaptive phase of CRISPR-mediated bacterial immunity, and perhaps in other analogous transposition reactions.

## Introduction

CRISPR-based bacterial immunity uses information from a prior infection stored in the chromosome to target and destroy invading viruses or plasmids. The adaptive phase of this defense system involves the deposition of short pieces of DNA (proto-spacers) from foreign nucleic acids into the chromosome as spacers—the storage units of immunological memory (1,2). The CRISPR locus has a [leader-(repeat-spacer-repeat)_n_] organization, and the long primary transcript derived from it is processed by Cas (CRISPR-associated) and/or host nucleases into mature CRISPR-RNAs (crRNAs) (3–6). The crRNAs, together with Cas effector protein(s), assemble the interference complex for recognition of invading nucleic acids through concerted protein-nucleic acid and base pairing interactions followed by their cleavage and destruction.

CRISPR-Cas systems are classified into two broad categories (7,8). Class 1 systems employ a multi-protein-RNA interference complex for target recognition, termed Cascade. Class 2 systems utilize RNA guided target recognition and cleavage by a single protein, i.e., Cas9 or Cas12. These two classes are further divided into several types and subtypes (6,9,10). Self-targeting is avoided via multiple mechanisms, including recognition of a proto-spacer adjacent motif (PAM) immediately bordering the crRNA-complementary DNA sequence (8,11–16).

Although CRISPR-Cas systems differ markedly in the execution of immune defense, the adaptation phase (invading DNA ➔ pre-spacer ➔ proto-spacer ➔ integrated spacer) is well conserved but still incompletely understood (17,18). The conserved Cas1 and Cas2 proteins are essential (and sufficient in some systems) to carry out spacer acquisition *in vivo* (19,20). In the *Escherichia. coli* (Type I-E) and *Enterococcus. faecalis* (Type II-A) Cas1-Cas2 crystal structures, a pair of Cas1 dimers flank a central Cas2 dimer on either side in a butterfly arrangement, providing a ridge that can accommodate the pre-spacer DNA (that includes the PAM sequence), or the proto-spacer DNA (that lacks a complete PAM), with splayed out single strand termini (21–23). The *E. faecalis* complex reveals a second ridge that can be occupied by the acceptor DNA, suggesting its integration competence (23). Other Cas proteins such as Cas4, Csn2 and Cascade/Cas9 may improve spacer acquisition efficiency, while also contributing to the mechanism for discrimination of non-self (pre-spacer) from self (integrated spacer) (24,25). In addition, bacterial nucleoid associated proteins such as IHF (integration host factor) assist the integration reaction and enhance target specificity in the *E. coli* Type I-E system (26). Proto-spacer generation in the Type I and II systems involves PAM recognition within the prespacer, but also the exclusion of a complete PAM from the proto-spacer to be integrated (24,25).

Proto-spacer integration is mechanistically analogous to DNA transposition, with terminal 3’-hydroxyls attacking the target phosphodiester bonds at the leader-repeat (L-R) junction on one strand and the repeat-spacer (R-S) junction on the other (19) (Figure 1A). These transesterification reactions are carried out by the Cas1 active sites of the Cas1-Cas2 complex. Repair of the intermediate containing the [proto-spacer]-[repeat] strand fusions completes the reaction with a duplication of the repeat sequence, one copy each on either side of the spacer insert (Figure 1A). The nucleophilic attacks by the 3’-hydroxyls are thought to occur in a stepwise fashion with bending of DNA to facilitate the reaction (23).

**Figure 1.**
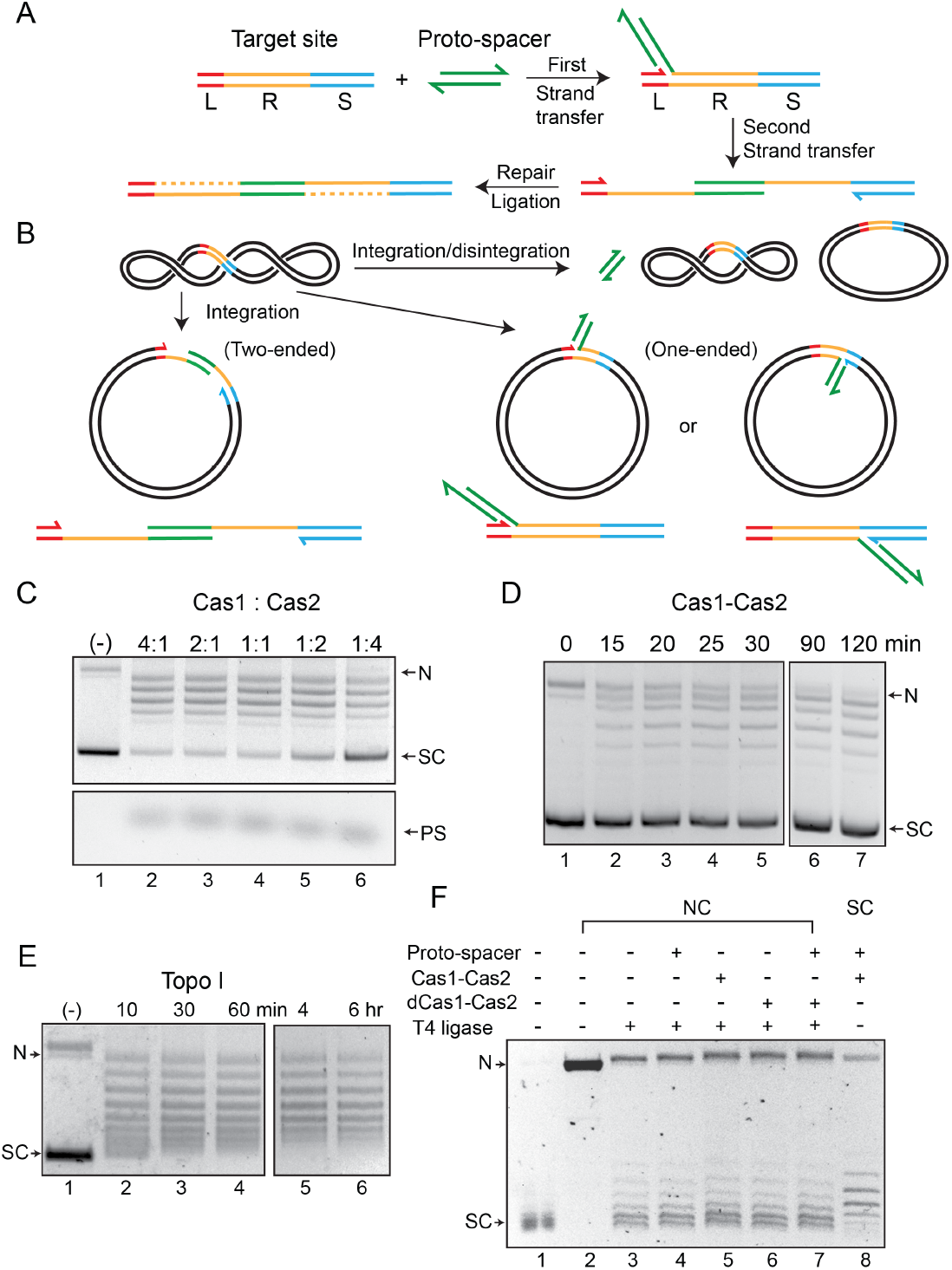
Cas1-Cas2 integrates and disintegrates proto-spacers at the CRISPR locus in a supercoiled plasmid. **A**. Illustration of protospacer (PS, green) integration into a minimal CRISPR locus comprised of the leader (L, red), repeat (R, yellow) and spacer (S, blue). The reaction follows the cut- and-paste DNA transposition mechanism with a duplication of the repeat sequence. The 3’-hydroxyls that perform nucleophilic attacks at the L-R junction (top strand) and the R-S junction (bottom strand) are indicated by the split arrowheads. **B**. In vitro reactions were performed with a supercoiled acceptor plasmid containing the minimal CRISPR locus (L = 11 bp; R = 36 bp; S = 27 bp). Each strand of the proto-spacer was 26 nt long, with four single stranded 3’-proximal nucleotides. The products of two-ended (complete) and one-ended (partial) proto-spacer integration and of protospacer integration-disintegration are diagrammed. Plasmid supercoiling will be reduced if the free DNA end undergoes rotation between integration and disintegration. **C**. The target plasmid and the proto-spacer were reacted with Cas1-Cas2 mixtures in the indicated molar ratios. After 2 hr incubation, reactions were analyzed by agarose gel electrophoresis and ethidium bromide staining. A split view of the top and bottom portions of the gel is presented to show the plasmid and proto-spacer bands. **D**. Reactions were similar to those shown in C, and utilized a Cas1 to Cas2 molar ratio of 2:1. **E**. The substrate plasmid used for the reactions in **C** and **D** was treated with a sub-optimal amount of E. coli topoisomerase I to follow the pattern of DNA relaxation over time. **F**. The T4 ligase reactions (lanes 3-7) were performed on the plasmid containing the L-R-S target site nicked with Nb BtsI (lane 2). The Cas1-Cas2 integration-disintegration reaction (lane 8) utilized the same plasmid in its supercoiled form (lane 1). Reactions were analyzed by gel electrophoresis in the presence of chloroquine (0.4 μg /ml). The DNA bands from the T4 ligase (lanes 3-7) and Cas1-Cas2 (lane 8) reactions are offset by one-half twist/writhe, as indicated by their staggered patterns. SC = supercoiled; N = nicked; PS = proto-spacer.

Crystal structures of Cas1, Cas2 proteins and of Cas1-Cas2 complexes (21–23,27,28), as well as cryo-electron microscopy (cryo-EM)structures of Cas1-Cas2-Csn2 complexes (29), are consistent with the transposition mechanism of proto-spacer integration. However, they also raise a number of intriguing questions. Are the adaptation steps--pre-spacer recognition, its processing into proto-spacer, and proto-spacer insertion into the CRISPR locus—carried out by the same complex, perhaps in coordination with bacterial helicase/nuclease proteins (17,30–32)? Or, are processing and insertion performed by separate complexes in spatially and/or temporally distinct steps? Does *in vitro* integration of a pre-processed proto-spacer by Cas1-Cas2 fully recapitulate the features of the *in vivo* reaction?

Here, we probe the mechanism of *S. pyogenes* (Spy) Cas1-Cas2 catalyzed integration *in vitro.* We show that partial (one-ended) integration is the most common reaction product. Attempts by Cas1-Cas2 to complete proto-spacer insertion by the integration of the second strand are most often accompanied by the reversal of integration (disintegration) of the first strand. As a result, complete integration of a proto-spacer into the target site is exceedingly rare. Nevertheless, the incomplete intermediates formed by disintegration may be rescued via repair pathways known to operate during DNA transposition. We conclude that disintegration is an intermediate step in at least one mode of integration, which may be accomplished by more than one pathway.

## Results

### Proto-spacer insertion into a nominal CRISPR locus present in a supercoiled plasmid

The *in vitro* insertion of a proto-spacer into a target site by Spy Cas1-Cas2 proteins (33) is inefficient, suggesting that the system may not fully represent the *in vivo* reaction. It may be limited by missing accessory components or because it is uncoupled from preceding DNA processing steps. We reexamined this reaction using a negatively supercoiled plasmid harboring the target site as ‘leader-repeat-spacer’ (Figure 1B).

The prominent outcome was the conversion of the supercoiled plasmid into a relaxed distribution (Figure 1C, D). The formation of a spacer integrant joined to the leader-proximal repeat end (L-R) on one strand and the spacer-proximal repeat end on the other (R-S) was barely detectable. Only a fraction of the product migrated as a nicked circle expected for a complete proto-spacer integration event or semi-integration events at either the L-R or R-S junction (Figure 1C, D). We confirmed the presence of the proto-spacer in the ‘nicked’ plasmid band by labeling it with Cy5 (Supplementary Figure S1A). By contrast, the Cy5-labeled proto-spacer was not inserted into the relaxed plasmid. Optimal reaction occurred at 4:1 and 2:1 molar ratios of Cas1: Cas2, with increasing Cas2 inhibiting the reaction mildly (Figure 1C). These results are consistent with previous biochemical and structural data suggesting Cas1_4_-Cas2_2_ as the active insertion complex, with Cas1 performing the catalytic steps (21,22) (Supplementary Figure S1B-D).

The predominance of the relaxed (closed circular) plasmid over the ‘nicked’ plasmid in the product indicates that the Cas1-Cas2 complex fails to complete proto-spacer integration in most cases. The major *in vitro* activity of Cas1-Cas2 is the reversal of semi-integration events (‘disintegration’) mediated by the 3’-hydroxyl exposed next to the integration junction.

### Topological features of the Cas1-Cas2 reaction in supercoiled plasmids

We employed topological analyses to address potential structural distortions of target DNA induced by Cas1-Cas2 during proto-spacer integration (21–23). We probed linking number changes (ΔLks) accompanying integration-disintegration in a supercoiled plasmid or strand closure in a nicked plasmid. Provided the Cas1-Cas2 induced topological stress within a target site is stable throughout the reaction, it will be preserved in the plasmid topoisomers formed by integration-disintegration (Figure 1B, C). In principle, the target topology in a Cas1-Cas2 bound nicked plasmid may also be captured by resealing the nick with DNA ligase.

DNA relaxation by Cas1-Cas2 occurred progressively, although not in a regular stepwise fashion (Figure 1D). The disappearance with time of the faster migrating bands formed early in the reaction, and the corresponding increase in the more relaxed distribution, is consistent with more than one ‘semi-integration-disintegration’ event occurring in the same plasmid molecule. The relaxation pattern was distinct from a topoisomerase I-generated ladder (ΔLk = +1) (Figure 1E), suggesting that the broken strand goes through multiple rotations before it is resealed.

Integration-disintegration by Cas1-Cas2 and nick closure by T4 DNA ligase resulted in similar topoisomer distributions (Supplementary Figure S1E). The slight shift in the center of the distribution from the Cas1-Cas2 reaction is consistent with supercoiling strain (writhe or twist) being sequestered by Cas1-Cas2 interaction with DNA. However, the possibility that relaxation by Cas1-Cas2 had not reached equilibrium cannot be ruled out. The relaxed plasmid topoisomers formed by ligation in the absence of Cas1-Cas2 reflect differences in twist or writhe in individual DNA molecules due to thermal fluctuations. As the gel fractionation did not include an intercalating agent, the position of a band below the slowest migrating fully relaxed form (ΔLk = 0) indicates only its absolute deviation from the relaxed state. A molecule with ΔLk = +1 (positively supercoiled) and one with ΔLk = −1 (negatively supercoiled) would show the same migration.

Next, we analyzed the integration-disintegration and nick ligation reactions by separating DNA topoisomers in the presence of chloroquine. Unwinding of the double helix by the intercalator introduces compensatory positive supercoils in covalently closed circular molecules. Each topoisomer is separated from its nearest neighbors (migrating above and below) by one supercoil, the faster migrating one being more positively supercoiled. Nearly identical topoisomer distributions were obtained when the nicked plasmid was ligated in the absence of Cas1-Cas2 (Figure 1F; lane 3) or in the presence of either Cas1-Cas2 (Figure 1F; lanes 5) or the catalytically inactive dCas1-Cas2 (Figure 1F; lane 6). Addition of protospacer by itself (Figure 1F; lane 4) or together with dCas1-Cas2 (Figure 1F; lane 7) to the ligase reaction did not alter this distribution. The topoisomer distribution of the Cas1-Cas2 plus protospacer treated supercoiled plasmid (Figure 1F; lane 8) was centered at roughly one more negative supercoil than that obtained from nick closure in the presence of dCas1-Cas2 and proto-spacer (Figure 1F; lane 7) (Supplementary Figure S2).

The above results suggest that Cas1-Cas2 either stably constrains local negative supercoiling when bound to a target site in a supercoiled plasmid (without, however, impeding free rotation of the strand nicked by Cas1), or has had insufficient time to effect complete plasmid relaxation. For a target site present in a nicked plasmid, the Cas1 - Cas2 induced topological stress is presumably dissipated via the nick before being sealed off by ligation. The observed ΔLk of ~1 between the Cas1-Cas2 relaxed and T4 ligase sealed plasmid distributions (Figure 1F; Supplementary Figure S2) is consistent with the unwinding of one DNA turn within the target sequence during proto-spacer integration. This torsional strain may promote the formation of the repair-ready intermediate in which single stranded repeat sequences flank the inserted proto-spacer (Figure 1A).

### Proto-spacer insertion into target sites in linear DNA

Previous studies demonstrated semi-integration of a proto-spacer into a linear DNA target, and indirectly suggested full integration (33). The effect of DNA topology on the disintegration reaction could account for the apparent difference in the activities of supercoiled and linear targets in integration. In order to characterize the integration steps in further detail, we turned to reactions with linear substrates (Figure 2A).

**Figure 2.**
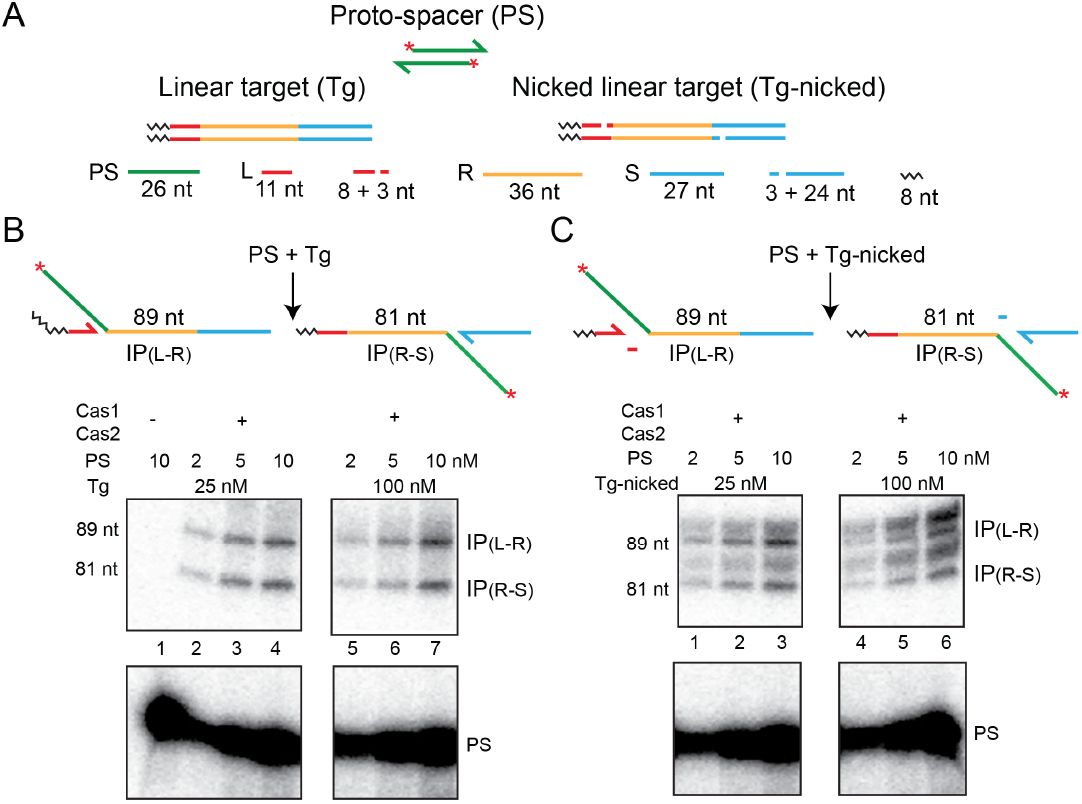
Cas1-Cas2 integrates proto-spacers into linear DNA targets. **A**. In the schematics, the proto-spacer (32P-labeled at the 5’-end; asterisk) and the linear target sites (intact or containing strand nicks) are color coded as in Figure 1. The nicks were placed three nts from the repeat, within the leader and the spacer on the top and bottom strands, respectively. A short oligonucleotide sequence adjacent to the leader (but unrelated to it) is represented by the black wavy lines. **B**, **C**. The expected integration products and their sizes are shown at the top. Reactions were analyzed by electrophoresis in 10% denaturing polyacrylamide gels, and bands were visualized by phosphor-imaging. Only the portions of the gel containing the relevant products and the unreacted proto-spacer are shown here (see Supplementary Figure S3 for the uncropped gel image). IP = Integration product; PS = Proto-spacer.

Consistent with previous findings, we observed insertions of the 5’-end-labeled proto-spacer at the leader side (L-R junction) (89 nt band) or at the spacer side (R-S junction) (81 nt band) (Figure 2B). L-R and R-S insertions were approximately equally prevalent (1.1: 1), although L-R insertions were more abundant at very short time points. This is consistent with semi-integration events initiated at L-R being converted to full-integration events by R-S insertion, as was suggested previously (33). However, alternative explanations cannot be ruled out (see below). We noticed additional faint product bands that are likely due to off-target events (Supplementary Figure S3), but these do not change the interpretations of the main results.

In order to address whether the products of integration could be stabilized by preventing potential disintegration, we used target sites with nicks placed within the leader and the spacer 3 nt away from the normal integration sites (Figure 2A). The short 3 nt product (5’NNN-OH3’) formed during integration would not be stably hydrogen bonded to the complementary strand and is expected to diffuse away from the reaction center. The increased spacing of the 3’-hydroxyls of the shortened leader or spacer from the integration junction would block or curtail disintegration.

The nick-containing target yielded, in addition to the expected 81 nt and 89 nt bands, product bands migrating slightly higher than each (Figure 2C). The flexibility afforded by the nick resulted in integration at the phosphodiester positions immediately adjacent to the nick, increasing the product length by one and two nucleotides, respectively. The overall amount of the integration product was increased by ~3 fold in comparison to that formed from the target without a nick.

Although disintegration by the leader or spacer 3’-hydroxyl is reduced in the nicked target site, water-mediated disintegration could still be an impediment to stable integration (see below). The similar amounts of L-R and R-S insertion product bands may arise from equal and independent one-ended insertions in the same target or in separate targets, or from completion of two-ended insertions within individual targets. A third possibility is that, within an intermediate containing a semiintegrant, integration of the second strand is coupled to disintegration of the first. In this case, the one-ended products from L-R and R-S junctions would be equal at steady state. Their absolute amounts would be dictated by how strongly they are destabilized by disintegration under a given set of reaction conditions. The difference between nearly zero and readily detectable integration in supercoiled versus linear targets can be accounted for by this interpretation.

### Sequential and coordinated strand transfer during proto-spacer insertion by Cas1-Cas2

Current evidence suggests independent recognition of L-R and R-S junctions by Cas1-Cas2 but initiation of integration at L-R (33). We examined the order of strand transfer between the two junctions using modified proto-spacers and target sites.

The modified proto-spacers contained base insertions either in both strands (5’T3’/3’T5’)_n_ or in only one strand (5’T3’/-)_n_, with no additional changes from the native sequence (Figure 3A). As a result, the length of a modified strand was increased from the normal 26 nt by the number of inserted Ts. However, the 22 bp double stranded region and the 4 nt 3’-overhangs at either end were retained as in the native protospacer. In the modified target sites, (5’T3’/3’T5’)_n_ insertions replacing a central 16 bp segment of the repeat were flanked on either side by 10 bp each of the leader-proximal and spacer-proximal repeat sequences. Consequently, the repeat lengths varied from 25 nt to 30 nt and 35 nt on each strand in the individual insertion derivatives (compared to 36 nt fully complementary strands in the native target). Based on the results from prior studies (33,34), the L-R and R-S junctions of the modified target sites were expected to be integration competent. Furthermore, the premise that the inherent structural pliability of (T/T)_n_ or (T/-)_n_ would at least partly compensate for changes in the absolute lengths of the Cas1-Cas2 substrates was borne out by the experimental results (Figure 3B, C).

**Figure 3.**
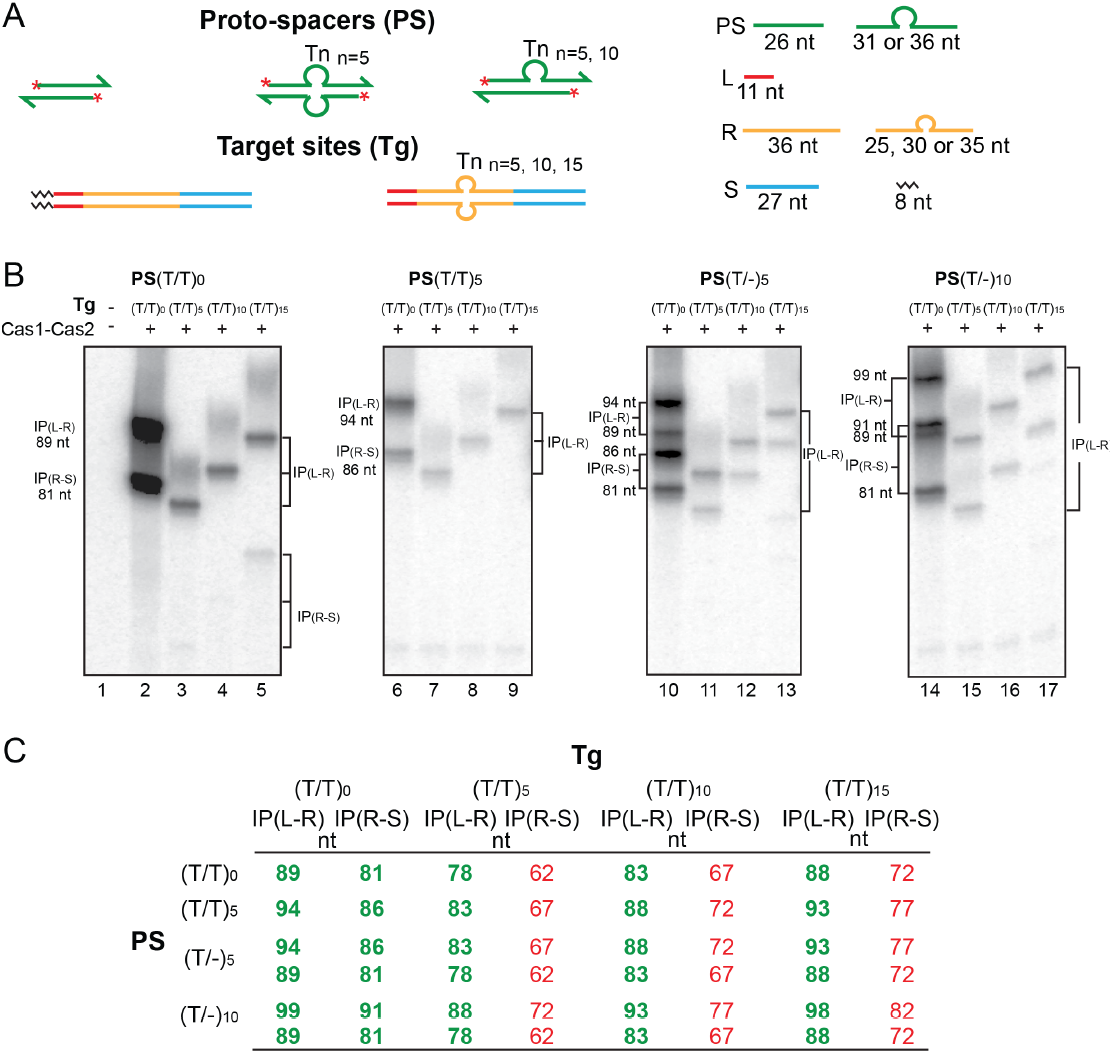
Cas1-Cas2 integrates modified proto-spacers into modified target sites. **A**. Schematics of the native and modified versions of the protospacers (labeled at the 5’-ends on both strands; asterisks) and target sites used for integration assays. **B**. The reactions were performed and analyzed as described under Figure 2. (see Supplementary Figure S4 for uncropped gel images). **C**. The expected integration products from the various protospacer and target site pairings are summarized. The (T/T)_0_ designation refers to the unmodified proto-spacer and target. The observed products from the reactions shown in **B** are in bold green font; those not detected or detected in extremely small amounts are in normal red font. IP = Integration product.

The native (T/T)_0_ proto-spacer as well as the ones carrying T-insertions, (T/T)_n_ or (T/-)_n_, gave the predicted integration products at the L-R and R-S junctions with the native (T/T)_0_ target site (Figure 3B; lanes 2, 6, 10 and 14). As the two strands of a (T/-)_n_ proto-spacer were unequal in length, their integration at each junction resulted in two products with different gel mobilities (four bands in lanes 10 and 14). For the (T/T)_5_, (T/T)_10_ and (T/T)_15_ targets, integration occurred almost exclusively at the L-R junction (Figure 3B; lanes 3-5, 7-9, 11-13 and 15-17). The corresponding products at the R-S junction were at or below the levels of detection (Figure 3B; Supplementary Figure S4). Extended phosphorimaging brought to light weak bands shorter than the predicted authentic integration products (Supplementary Figure S4). Although they are not directly relevant to our analysis, their possible origin is considered in Text S1 under Figure S4. Reactions of the modified proto-spacers with the modified target sites unveil a strict hierarchy in the choice of the L-R and R-S junctions during proto-spacer acquisition. Figure 3C differentiates the experimentally observed products from the undetected (or highly underrepresented) potential products. The strong L-R preference could occur at the level of target recognition (which then triggers catalysis) or at the level of catalysis (with no preferential recognition of either the L-R or the R-S junction). Structural snapshots of the *E. faecalis* Cas1-Cas2 integration complex suggests that, following a stochastic L-R/R-S search, preferential interaction with the L-R junction is established (22). Catalysis of the initial one-ended insertion at this target site follows. Within this intermediate, perhaps a ruler-like mechanism is used to promote R-S junction interactions that culminate in the second strand transfer. When the search fails to identify an appropriately positioned R-S, as would be the case with the modified targets, the reaction is limited almost entirely to the L-R junction.

### Proto-spacer insertion into half-target sites

One question raised by the results from the modified target substrates is whether the [L-R]-before-[R-S] rule in proto-spacer strand transfer is manifested only when both these junction sequences are present in *cis* within a target site. Previous mutational analysis suggests that either the L-R or the R-S junction is used equally well for integration when only one of them is functional (21,22). We now assayed the integration of a proto-spacer into linear L-R and R-S half-target sites to test R-S functionality in the absence or presence of L-R.

In reactions with the individual L-R and R-S half-sites, insertion products were obtained with either half-site alone (Figure 4; lanes 1-14). Consistent with the independent recognition of the L-R and R-S junctions by the Cas1-Cas2 complex, an R-S half-site was codominant with the ‘minimal’ L-R half-site (containing 18 bp of the repeat) when both were present in the same reaction (Figure 4; lanes 15-18). However, extension of the repeat segment in the minimal L-R halfsite by 4 bp, L-R(+4), restored L-R dominance to the same degree as that displayed by the modified full linear targets (Figure 4; lanes 19-22). While the insertion product was formed from L-R(+4), none was detected from either the minimal R-S half-site (containing 18 bp of the repeat) or the R-S(+6) half-site (containing additional 6 bp of the repeat).

**Figure 4.**
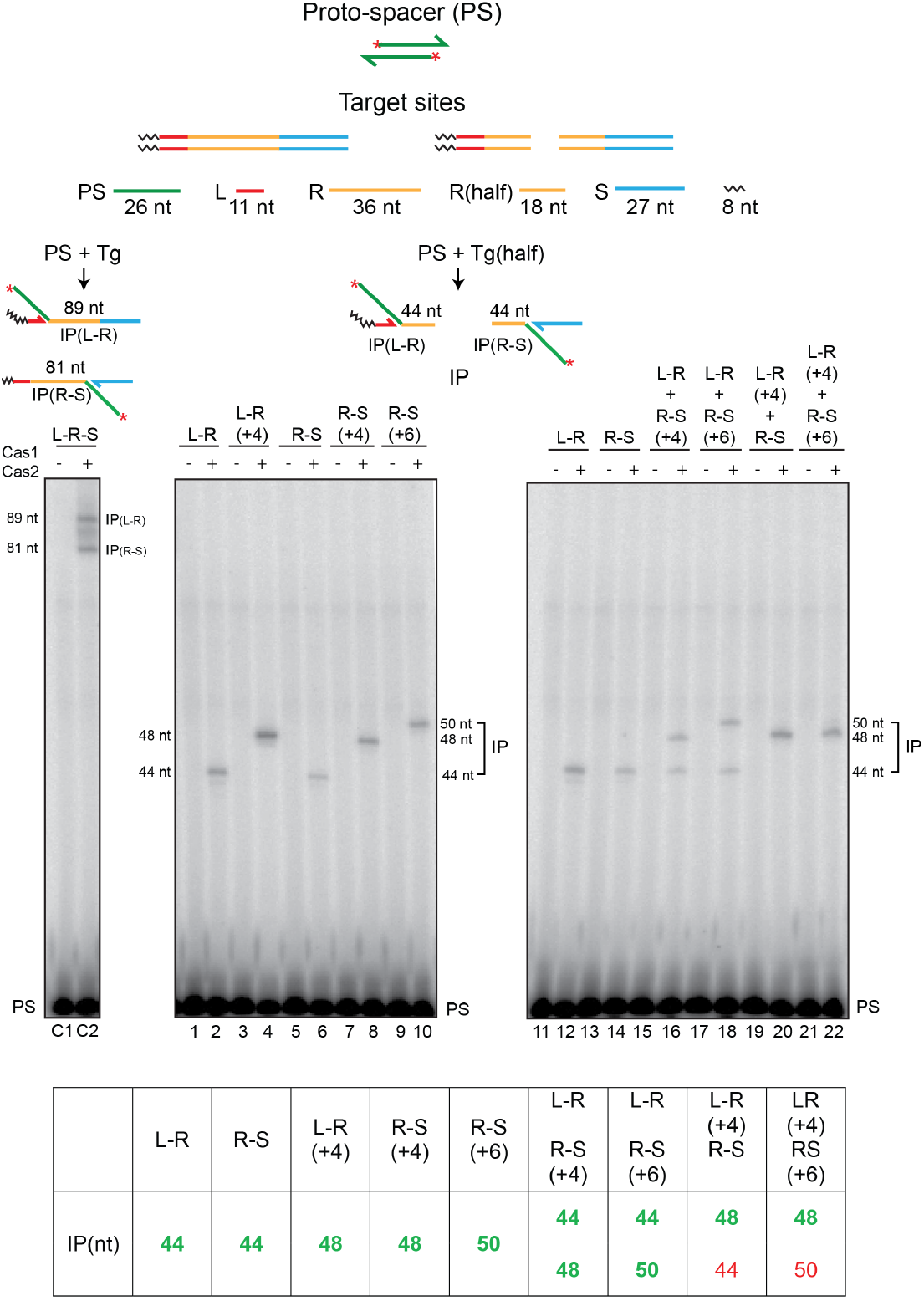
Cas1-Cas2 transfers the proto-spacer into linear half-target sites. The proto-spacer labeled with 32P at its 5’-ends and the linear full- and half-target sites are diagrammed at the top. The minimal L-R or R-S half-site contained 18 bp of the L-proximal or R-proximal repeat sequence, respectively. Each half-site is the product of splitting a full-site into two exactly at the center of the 36 bp repeat. Reactions contained the labeled proto-spacer together with the indicated minimal L-R or R-S half-sites, or half-sites containing longer repeat sequences (+4 or +6 bp), as well as combinations thereof. Reactions were analyzed as described under Figures 2 and 3. The lanes C1 and C2 denote the control reaction with a normal target site containing the full-length repeat. The predicted integration products from the individual and mixed half-target site reactions are listed at the bottom. The experimentally observed products are distinguished from the undetected ones by the bold green and normal red fonts, respectively. IP = integration product; PS = proto-spacer.

Thus, the hierarchical dominance of L-R seen in the full-target sites (Figure 3B) is lost when R-S is physically unlinked from the minimal L-R. However, L-R(+4) is dominant over R-S or R-S(+6) even in the absence of linkage. The extent of repeat sequence present adjacent to the L-R junction and/or specific repeat sequences distal to this junction may contribute to the priority in target site recognition and order of strand transfer during spacer acquisition. It is possible that the integration reaction occurs within a Cas1-Cas2 complex harboring a non-covalently dimerized pair of half-sites, (L-R)2; (R-S)2 or (L-R)-(R-S), and the L-R(+4) dominance is manifested in this context.

### Apparent integration-disintegration antagonism during spacer acquisition

‘Cut-and-paste’ transposition during integration involves an unusually short transposon (the proto-spacer) and a particularly long target site (the CRISPR repeat). Their nearly matched sizes may partly relieve the steric impediments to insertion posed by DNA persistence length. Nevertheless, the strain within the strand transfer intermediate could potentially trigger disintegration mediated by the 3’-hydroxyl exposed at the leader or spacer end. Disintegration may offer a protective mechanism to prevent off-target integration events that could potentially imperil the integrity of the bacterial genome (33). The conformational dynamics of authentic integration at the CRISPR locus may displace the 3’-hydroxyl from the proto-spacer-repeat junction or misorient it with respect to the scissile phosphate. We investigated how disintegration might influence the second step of proto-spacer integration by taking advantage of substrates that mimic a semi-integrated spacer.

The reactions were carried out in substrates assembled by hybridizing four oligonucleotides to represent semi-integration at the L-R junction (Figure 5A). The 3’-hydroxyl of the unintegrated proto-spacer strand would serve as the nucleophile for the second integration step. Similarly, the 3’-hydroxyl of the ‘leader strand’, and potentially water, could provide the nucleophiles for the disintegration reaction. When the 5’-end of the unintegrated proto-spacer strand was labeled, integration was observed at the R-S junction (81 nt band) (Figure 5A; lanes 2-6). The reaction was rapid, and was nearly saturated within a minute under the conditions employed. Along with the 81 nt product, a smaller amount of the companion 89 nt product was also formed, whose yield showed a linear increase with time (Figure 5A; lanes 2-6). Formation of this minor product suggests that the phosphodiester bond at the L-R junction was restored by 3’-hydroxyl-mediated disintegration in at least a subpopulation of the starting substrate molecules. This junction could then serve as the target site for strand insertion from a disintegrated protospacer. Consistent with this interpretation, labeling the 5’-end of the leader strand revealed the 92 nt product expected for disintegration carried out by the leader 3’-hydroxyl to re-form the L-R junction (Figure 5A; lanes 2’-6’). Strikingly, disintegration was more prominent than integration.

**Figure 5.**
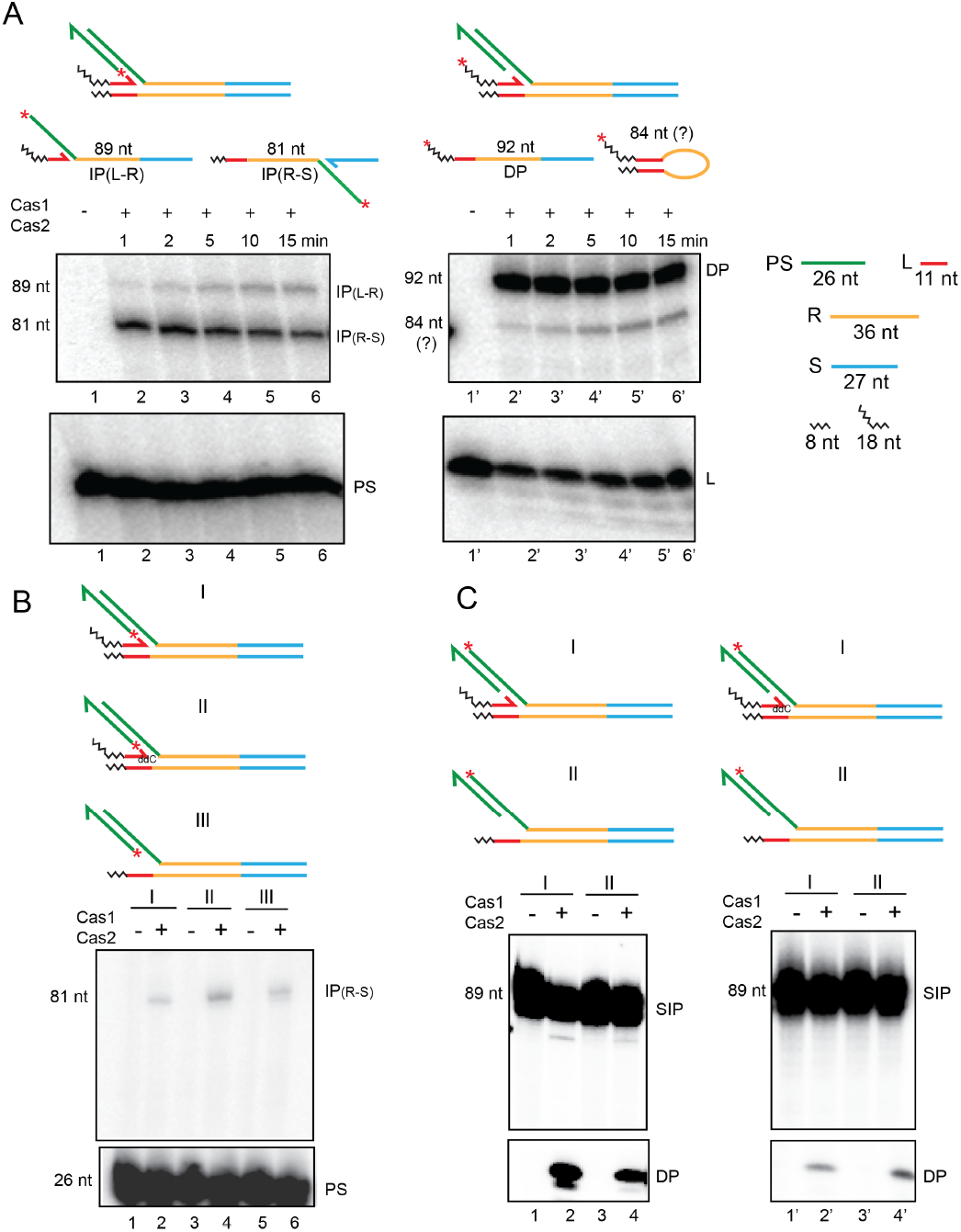
Second integration and disintegration occur in substrates mimicking a semi-integrated proto-spacer. **A**. Schematic of semiintegrated protospacers at the L-R junction (top row) and the possible products formed from them by Cas1-Cas2 action (bottom row). The 32P-label was placed at the 5’-end of the unintegrated proto-spacer strand or the leader strand to follow the second integration and disintegration events, respectively. **B**. Integration of the 32P-labeled proto-spacer strand at the RS junction was followed in the indicated substrates. **C**. Disintegration was assayed in the same substrates as in **B**, except that the 32P-label was placed at the 5’-end of the integrated proto-spacer strand. ddC = dideoxy C; SIP = Semi-integrated proto-spacer; IP = integration product; DP = disintegration product; PS = proto-spacer.

An unexpected product that migrates slightly slower than the 81 nt band was observed in reactions containing the 5’-end label on the leader strand (Figure 5A; lanes 2’-6’). This product, which showed a steady increase with time, suggests the potential utilization of the leader 3’-hydroxyl by Cas1-Cas2 in an integration-like reaction (Supplementary Figure S5 and Text S2). This uncharacterized product has no direct bearing on the possible functional link between integration and disintegration (see below and under ‘Discussion’).

In a substrate mimicking semi-integration at the R-S junction, Cas1-Cas2-promoted the second integration step at the L-R junction, yielding the expected 89 nt product (Supplementary Figure S5). Resealing of the R-S junction via disintegration mediated by the 3’-spacer hydroxyl was also evident from the smaller amounts of the 81 nt product formed by the semi-integration of a released protospacer at R-S. Unlike the first strand transfer step, which obeys a strict order during proto-spacer insertion, the second strand transfer may proceed from L-R to R-S or R-S to L-R with roughly equal efficiency. The prominence of disintegration and the kinetic correspondence between disintegration and the second integration reactions (Figure 5A) suggest that the two events are potentially coupled.

### Water-mediated disintegration of a semi-integrated proto-spacer

Disintegration from one of the semi-insertion sites (the L-R or the R-S junction) could produce a proto-spacer that is either free in solution or isconcomitantly inserted at the other site. The coupling of integration to disintegration, consistent with their reaction kinetics (Figure 5A), could be promoted by the conformational strain associated with orienting the reactive 3’-hydroxyl for the second transfer step. We wished to test whether preventing the disintegration reaction at the L-R junction in the semi-integrated state would reduce integration at the R-S junction.

We attempted to block the disintegration competence of the leader strand in the substrate with the proto-spacer semi-integrated at the L-R junction by placing a terminal 3’-dideoxy nucleotide, or by removing this strand altogether. Neither of these substrate designs eliminated or reduced the extent of integration at the R-S junction (81 nt band in Figure 5B). However, labeling the 5’-end of the proto-spacer strand integrated at the L-R junction revealed that disintegration occurred even in the absence of the leader strand (Figure 5C).

Water-mediated disintegration in the semi-integrated substrate could potentially assist the integration of the second proto-spacer strand in a coupled fashion. Disintegration efficiency of water alone (in the absence of the leader strand) was ~40% of that of the 3’-hydroxyl and water combined (in the presence of the leader strand) (Figure 5C, compare lanes 2 and 4). When the leader strand carried a di-deoxy 3’-end, there was some reduction (to ~70%) in water-mediated disintegration (Figure 5C; compare lanes 2’ and 4’), presumably due to steric interference. Such interference could be mitigated if, during a standard reaction, proto-spacer integration at the L-R junction is accompanied by the displacement or misalignment of the leader 3’-hydroxyl. We note that water is ~50% as competent a nucleophile as the 3’-hydroxyl for disintegration.

The 3’-hydroxyl-mediated and water-mediated disintegration reactions differ in that the former rejoins L to R (or R to S) whereas the latter leaves open the strand nick between L and R (or R and S). Processing of such a nick by DNA repair enzymes has functional implications in proto-spacer acquisition (see Discussion).

### Coupling between integration of the second strand and disintegration of the first

To better appreciate the extent of coupling between integration and disintegration, we modified the design of the semi-integrated state. The rationale was to join the integrated and unintegrated proto-spacer strands, so that the fate of both could be followed simultaneously.

In the ‘trombone’ substrates for testing coupled integration-disintegration (Figure 6), the proto-spacer strand was integrated at the L-R junction. Furthermore, the 3’ end of the spacer top strand was tethered to the 5’-end of the unintegrated proto-spacer strand. Our reasoning that the 30 nt long flexible single stranded tether (5’T_4_(CT)_10_T_6_3’; ~240Å) should have no (or only minimal) effect on Cas1-Cas2 activity was upheld by experiments. The spacer in these substrates was truncated to contain only 10 bp abutting the repeat. The shortened spacer had no adverse effect on strand transfer of an untethered proto-spacer from the L-R to the R-S junction (Figure 6A; lanes 2 and 4).

**Figure 6.**
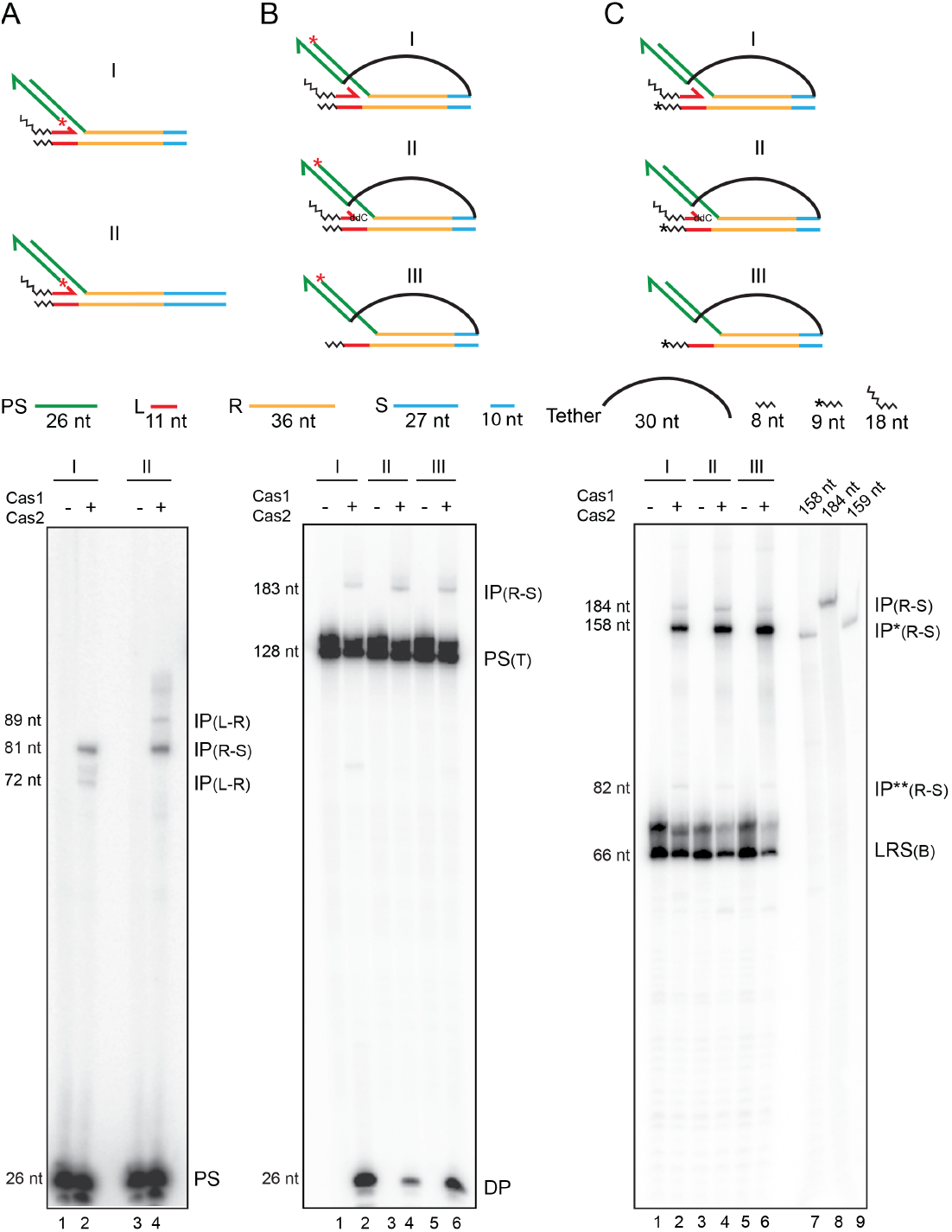
Second strand integration by Cas1-Cas2 occurs in concert with first strand disintegration. Schematics of the substrates used for the individual sets of assays (**A-C**) are placed above the respective gel panels. **A**. Integration reactions were performed in the two substrates that differed only in their spacer lengths. The truncated spacer (10 bp long) lacked the 17 bp distal to the R-S junction. **B**, **C**. The substrate configurations were similar for the two reaction sets, except for the position of the 32P label. In **B**, the integrated proto-spacer strand carried the label at the 5’ end (red asterisk). In **C**, one labeled nucleotide (shown by the asterisk) was added to the 3’ end of the bottom strand. IP = integration product; IP* = integration product coupled to disintegration; IP** = integration product uncoupled from (but preceded by) disintegration; DP = disintegration product; PS = protospacer; PS (T) = proto-spacer (tethered); LRS (B) = substrate bottom strand. Lanes 7-9 in **C**, oligonucleotide length markers.

The predicted products of the second integration event in a trombone substrate would differ by 26 nt (the size of the integrated protospacer strand), depending on whether it is associated with disintegration or not (Supplementary Figure S6). Labeling the 5’-end of the integrated proto-spacer strand would only reveal integration at the R-S junction without disintegration at the L-R junction (183 nt; Figure 6B) (Supplementary Figure S6A). Formation of this product was weak, though detectable, in all three substrates tested, two of which contained a leader strand (one with a hydroxyl and the other with a deoxy 3’-end) and the third lacked this strand (Figure 6B; lanes 2, 4 and 6). At the same time, disintegration mediated by the leader 3’-hydroxyl or by water was readily observed (26 nt; Figure 6B; lanes 2, 4 and 6) (Supplementary Figure S6A). Labeling the 3’-end of the bottom strand containing the R-S junction target site (by adding one α-^32^P-labeled nucleotide in a Klenow polymerase reaction) would reveal integration events coupled to disintegration (158 nt; Figure 6C) or uncoupled from it (184 nt; Figure 6C) (Supplementary Figure S6B). In all three substrates, the shorter product exceeded the longer one (Figure 6C; lanes 2, 4 and 6), by 8- to 10-fold when the leader strand carried a 3’-hydroxyl (Figure 6C; lane 2) or had a deoxy 3’-end (Figure 6C; lane 4), and by >15-fold when the leader strand was absent (Figure 6C; lane 6). The maximum yield of the 158 nt product in the absence of a leader strand suggests that the second integration is more efficient when disintegration is performed by water rather than by the 3’-hydroxyl.

Several features of the proto-spacer strand transfer steps are revealed by the trombone substrates (see also Supplementary Figure S6). First, integration of the second strand without disintegration of the first is quite infrequent. Second, disintegration of a proto-spacer from the L-R junction followed by its integration at the R-S junction in a two-step process is also rare. The IP(R-S)** band (82 nt; Figure 6C) is the result of such a reaction, namely, insertion of the 26 nt proto-spacer strand released from the L-R junction into the R-S junction. Thus, integration of the second strand and disintegration of the first are most often concerted events, and two-ended insertion events by the Cas1-Cas2 complex seldom occur *in vitro.* Finally, water being better at promoting disintegration coupled integration than the 3’-hydroxyl is significant. The strand nick left unsealed during the water-mediated reaction would permit seemingly abortive intermediates to be processed by bacterial DNA repair systems into successful spacer insertion (see ‘Discussion’). Thus, paradoxically, disintegration may be an important (but unappreciated) contributing factor in the adaption stage of CRISPR immunity.

## Discussion

Cas1-Cas2-mediated adaptation is strongly conserved across CRISPR systems, but is less understood than the much more diverse interference mechanisms (5,18,20,35,36). Our analysis indicates that the integration by Cas1-Cas2 *in vitro* starts with the insertion of one proto-spacer strand at the L-R junction and utilizes a ruler-like mechanism to attempt integration of the second strand at the R-S junction. However, a successful second integration event is most often associated with disintegration of the first one. We propose below plausible models for disintegration-promoted spacer acquisition that are accommodated by well-established DNA transposition mechanisms.

Two-ended proto-spacer integration, without intervening disintegration, is conceivable *in vivo*. A high-order adaptation complex in which Cas1-Cas2 is assisted by additional Cas proteins (e.g., Cas2, Csn2, Cascade/Cas9) and by bacterial factors (e.g., RecBCD, AddAb) (17,19,20,24,25,31) may deter disintegration by coupling DNA processing to strand transfer without proto-spacer dissociation. Dynamic remodeling of the complex in conformation and/or subunit stoichiometry, as suggested by Cas1-Cas2-Csn2 cryo-EM structures (29), could facilitate the individual reaction steps. However, if a pre-processed protospacer were to escape from the complex, it is still able to complete Cas1-Cas2-catalyzed, and disintegration-promoted, integration.

### From invading DNA to spacer deposition at the CRISPR locus: variations of a shared DNA transposition theme

An invading virus/plasmid DNA is processed by bacterial nuclease/helicase complexes into smaller sized fragments (pre-spacers), to be further processed by Cas proteins into proto-spacers suitable for integration at the CRISPR locus (17,31,32,37–41). The early and subsequent processing events may be coordinated by communication between the Cas proteins and nuclease/helicase proteins. In principle, the action of Cas1-Cas2 (perhaps with the assistance of partner Cas proteins) directly on the invading foreign DNA, or on shorter processed fragments generated from it, may give rise to unexcised proto-spacers with cleaved single stranded ends or fully excised free-standing proto-spacers, both of which would be competent to initiate the integration reaction (Figure 7). These situations would be analogous to replicative and cut-and-paste DNA transposition, respectively (42–44). In the cut-and-paste mechanism, only limited gap repair DNA synthesis is required to reestablish the native organization of the CRISPR locus, but with the newly acquired spacer (Figure 7). In the replicative mechanism, DNA synthesis is more extensive, and the intermediate is resolved via recombination to yield the same end product as in cut-and-paste transposition (Figure 7). Variations of these themes, combining aspects of replicative and cut-and-paste transpositions may also be envisaged (Supplementary Figure S7). Furthermore, the strand cleavage and strand transfer steps of proto-spacer insertion must necessarily engender safeguards against self-targeting of the inserted spacer as well as nonfunctional spacer orientation (Supplementary Figure S8; Text S3).

**Figure 7.**
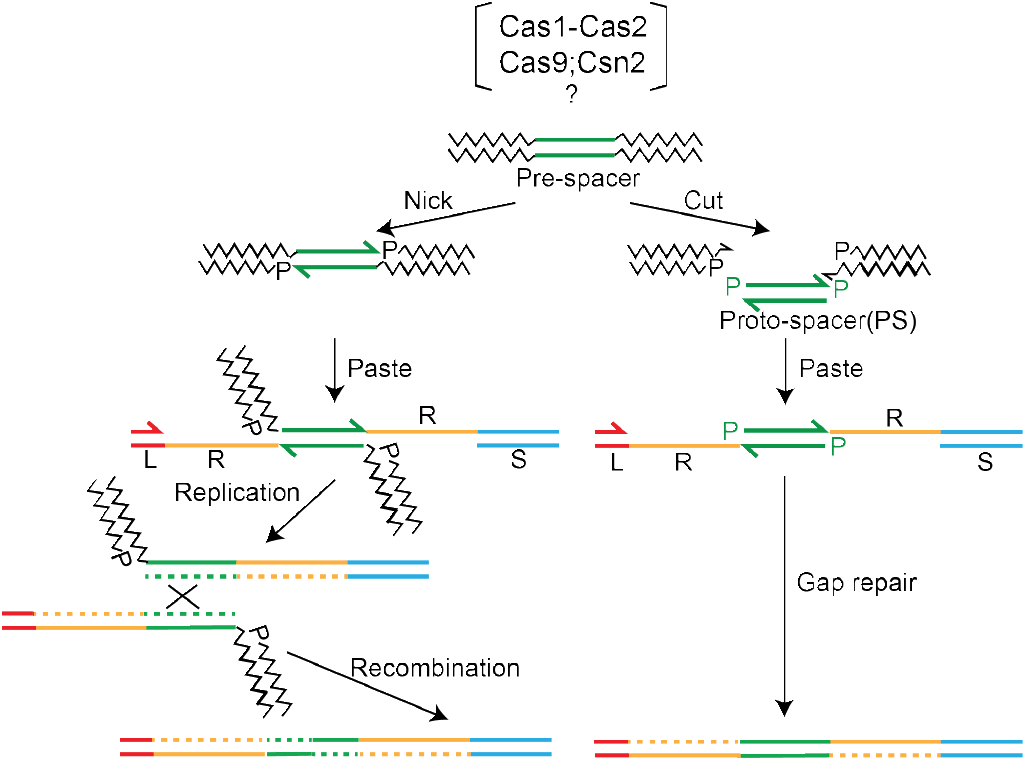
Two-ended integration of the proto-spacer by the spacer acquisition complex. Two possible schemes for proto-spacer integration are outlined. The pathway shown at the left is equivalent to replicative DNA transposition. The pre-spacer is nicked at the proto-spacer 3’ ends, which are then transferred to the leader-repeat (L-R) and repeat-spacer (R-S) junctions of the target site. Replication across L and PS (proto-spacer) followed by recombination within the duplicated PS completes integration. An alternative mode of processing the strand transfer intermediate involves removal of the pre-spacer DNA flanking PS, and gap repair by limited DNA synthesis and ligation (Supplementary Figure S7). The integration pathway shown at the right follows cut-and-paste DNA transposition. The steps involve excision of PS from the pre-spacer, strand transfer to L-R and R-S junctions, gap repair and ligation.

A recent single molecule FRET analysis suggests that the post-strand transfer complex (or post-synaptic complex; PSC) formed during proto-spacer integration may be resolved by transcription from the CRISPR leader accompanied by limited DNA replication (45). Resolution may also be achieved by replication, although it is argued to be less likely. The lag time in the arrival of a replication fork at the PSC in a slow-growing bacterium may cause disintegration of the proto-spacer and its potential reintegration in the non-functional orientation. Replication through the unwound repeat and proto-spacer would produce undesirable double strand DNA breaks. However, if the integration complex functions as a strand transfer-DNA repair machinery, the DNA ends may remain protected within it to be resolved rapidly (Figure 7; pathway at the left). Precedent exists for the involvement of the replisome in the repair of the strand transfer intermediate formed during the insertion of infecting phage Mu DNA into the *E. coli* chromosome (46).

### Cas1-Cas2-mediated integration of a pre-processed proto-spacer: assistance from disintegration

If ‘normal’ (two-ended) integration by the adaptation complex (Figure 1A; Figure 7) is disrupted after only one proto-spacer strand has been transferred to the target, salvage pathways may promote the maturation of such intermediates into complete spacer insertions. These pathways may also come into play during the integration of a proto-spacer that is dissociated from the adaptation complex and is captured by the Cas1-Cas2 complex. Integration under this scenario, which mimics the *in vitro* system, is expected to be predominantly one-ended. We suggest that disintegration events, which become prominent during the transfer of the second strand, play a central role in the salvage mechanisms.

Water-mediated disintegration of the proto-spacer at the L-R junction during strand transfer to the R-S junction will produce an intermediate with a gap in the bottom strand and a gap plus nick in the top strand (Figure 8). An analogous intermediate with the gap plus nick in the bottom strand may also be formed (Figure 8). Here, a proto-spacer is first transferred to the R-S junction with concomitant 3’-hydroxyl-mediated disintegration from the L-R junction. This is followed by a subsequent integration in the reverse direction, with disintegration at the R-S junction promoted by water. The spacer insertion event can be completed by gap-filling/displacement DNA synthesis followed by ligation (Figure 8). Thus, disintegration reactions mediated by the 3’-hydroxyl and by water are both relevant to the rescue of semi-integrated proto-spacers into fully integrated spacers.

**Figure 8.**
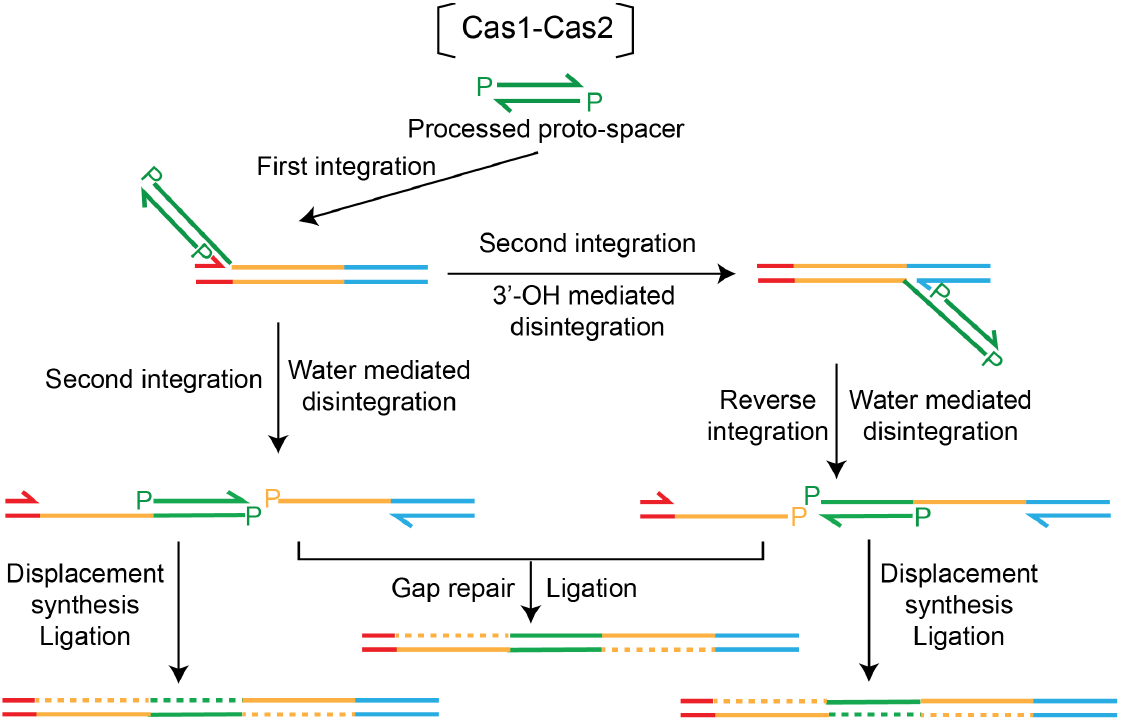
A model for disintegration-assisted integration. Integration of the proto-spacer by Cas1-Cas2 is predominantly one-ended. Transfer of the second proto-spacer strand is accompanied by concomitant disintegration mediated by the 3’-hydroxyl exposed at the L-R or R-S junction or by water. Water-mediated disintegration during the second strand transfer reaction preserves the nick between L and R or R and S that can be utilized by DNA repair machinery to complete PS insertion.

### A potential broader role for disintegration in DNA transposition

Disintegration is generally viewed as antithetical to the completion of the two strand transfer events during transposition reactions, which are analogous to proto-spacer insertion at the CRISPR locus. However, the *in vitro* Cas1-Cas2 reactions suggest that disintegration can generate DNA ends capable of triggering repair events that convert one-ended strand transfer intermediates into normal insertion products. This mechanism may be especially important when two-ended insertion is impeded by the transposon being considerably shorter than the persistence length of DNA. In a broader sense, disintegration may assist the movement of mini-transposons or short orphan transposons that are mobilized by ectopically supplied transposases. Disintegration need not always be counterproductive even during normal transposition, provided the DNA repair machinery is able to recommence the interrupted reaction. The Cas1-Cas2 system highlights the resourcefulness of biological catalysis in repurposing an apparently abortive reaction towards a physiologically meaningful outcome.

## Materials and Methods

### Plasmids

Protein expression plasmids and substrate plasmids used for *in vitro* assays were kindly provided by the Doudna laboratory (33).

### Oligonucleotides

Oligonucleotides used for the assembly of proto-spacers and target sites were purchased from IDT (listed in Supplementary Table S1). Oligonucleotides > 100 bp long were synthesized as two separate chains, and were ligated with the assistance of a short bridging splinter oligonucleotide. The 5’-end was made competent for ligation by phosphorylating it with ATP using T4 polynucleotide kinase.

Hybridizations were performed by mixing the requisite oligonucleotides in 20 mM HEPES-NaOH (pH 7.5), 25 mM KCl and 10 mM MgCl2, immersing the tube containing the mixture in a water bath at 90° C for 5 min, and letting the bath cool slowly to room temperature. After adding DTT (dithiothreitol) and DMSO (dimethyl sulfoxide) to final concentrations of 1 mM and 10%, respectively, they were immediately used for reactions or stored at −20° C until use (not more than a week).

### Proteins

His6-tagged Cas1, Cas2 and Csn2 were expressed individually as MBP (maltose binding protein) fusions in *E. coli*, and were purified by affinity chromatography over Ni-sepharose using standard protocols. When desired, MBP was removed by cleavage at a TEV protease target site placed at the fusion junction. MBP-free proteins were purified by Ni-affinity chromatography. Final protein preparations were dialyzed against 20 mM Tris-HCl (pH, 7.5), 150 mM KCl (Cas1; Csn2) or 500 mM KCl (Cas2), 1 mM TCEP (Tris(2-carboxyethy)phosphine hydrochloride), and stored at −70° C after addition of 10% glycerol (v/v). There was no difference in the activities between the MBP-fusion and MBP-cleaved forms of each protein. All data shown here were obtained using the fusion proteins.

### Radioactive labeling

5’-end labeling of oligonucleotides was carried out using γ-^32^P labeled ATP and T4 DNA ligase. For labeling the 3’-end, the Klenow fill-in reaction was performed in the presence of α-^32^P labeled dATP. A short-oligonucleotide with a 5’ T overhang hybridized to the oligonucleotide being labeled ensured the precise incorporation of one labeled A.

### Cas1-Cas2 activities

Reactions were performed in 20 mM HEPES-NaOH (pH, 7.5), 25 mM KCl, 10 mM Mg Cl_2_, I mM DTT and 10% DMSO (33) containing 5% PEG (polyethylene glycol 1500; Millipore Sigma). Incubations were carried out in 20 μl reaction mixtures at 37° C for 1 hr in most cases. In time course analyses, incubation periods ranged from 1 min to 1 hr, Reactions were stopped by addition of SDS (0.2% final; w/v) and heating to 95° C for 5 min. Samples were analyzed by electrophoresis in 1.5% agarose gels or 12% denaturing polyacrylamide-urea gels (19:1 crosslinking). In one set of assays probing plasmid topoisomer distributions, 0.4 μg chloroquine/ml was present in the agarose gel and the running buffer. DNA bands were visualized by ethidium bromide staining, fluorography or by phosphorimaging depending on the particular assays.

Reactions utilizing supercoiled plasmids contained ~10 nM plasmid DNA, ~67 nM Cas1 and ~33.5 nM Cas2 (2:1 molar ratio of Cas1: Cas2). In reactions containing a mixture of Cas1 and Cas1(H205A) (dCas1), the combined concentration of the two per reaction was kept as ~67 nM. For reactions utilizing ^32^P-labeled oligonucleotides, the DNA substrate was held at 25 nM with Cas1 and Cas2 at ~134 nM and 67 nM, respectively. In the Cas1-Cas2 cleavage assays (Supplementary Figure S8), Csn2 was included (where indicated) at a concentration of ~67 nM. Proteins were diluted from stock aliquots to a concentration of 1μM in the reaction buffer. Cas1-Cas2 or Cas1-Cas2-Csn2 were mixed in the requisite molar ratios and kept on ice for 5 min before addition to reaction mixtures.

### Topoisomerase reactions

A titrated amount of *E. coli* topoisomerase I was used to obtain partial relaxation of plasmid DNA. Reactions were carried out in the manufacturer recommended buffer in 30 μl volumes containing 300 ng supercoiled plasmid DNA per tube. After incubation at 37° C for varying times, reactions were quenched with SDS (0.2% final; w/v) and heated to 95° C for 3 min before electrophoresis in 1.5% agarose.

### Ligation of nicked plasmid DNA

Plasmid DNA, nicked at a single site using Nb.BtsI (New England Biolabs), was ligated using T4 DNA ligase and the ligation buffer (obtained from New England Biolabs). Each reaction mixture containing 300ng nicked plasmid and 200U ligase was incubated for 1 hr at 37° C before inactivating the enzyme by SDS addition (0.2% final; w/v) and heating to 95 C for 3 min. Samples were analyzed by electrophoresis in 1.5% agarose.

### Quantification of reaction efficiencies

^32^P-labeled bands on polyacrylamide gels were detected using a storage phosphor screen (Bio-Rad) at different exposure times to optimize signal detection without saturation of the screen. Scans were scanned in a Typhoon Trio phosphorimager (GE Healthcare), and image analysis was performed using the software Quantity One (Life Sciences Research) and ImageJ (NIH).

## Author Contributions

C.M. and K.J. performed research, I.J.F. and M.J. wrote the paper with input from all co-authors. The authors declare no conflict of interest.

This article contains supporting information online.

## Acknowledgments

We thank Drs. Addison Wright and Jennifer Doudna for Cas1-Cas2 plasmids and members of the Finkelstein and Jayaram labs for helpful discussions. The work was supported by NIGMS R01GM124141 (to I.J.F.), NSF MCB-1049925 and MCB-1949821 (to M.J.) and the Welch Foundation grants F-1274 (to M.J.) and F-1808 (to I.J.F.).

