## Supplemental File for "Disintegration promotes proto-spacer integration by the Cas1-Cas2 complex"

### SUPPLEMENTARY DATA

Figure S1

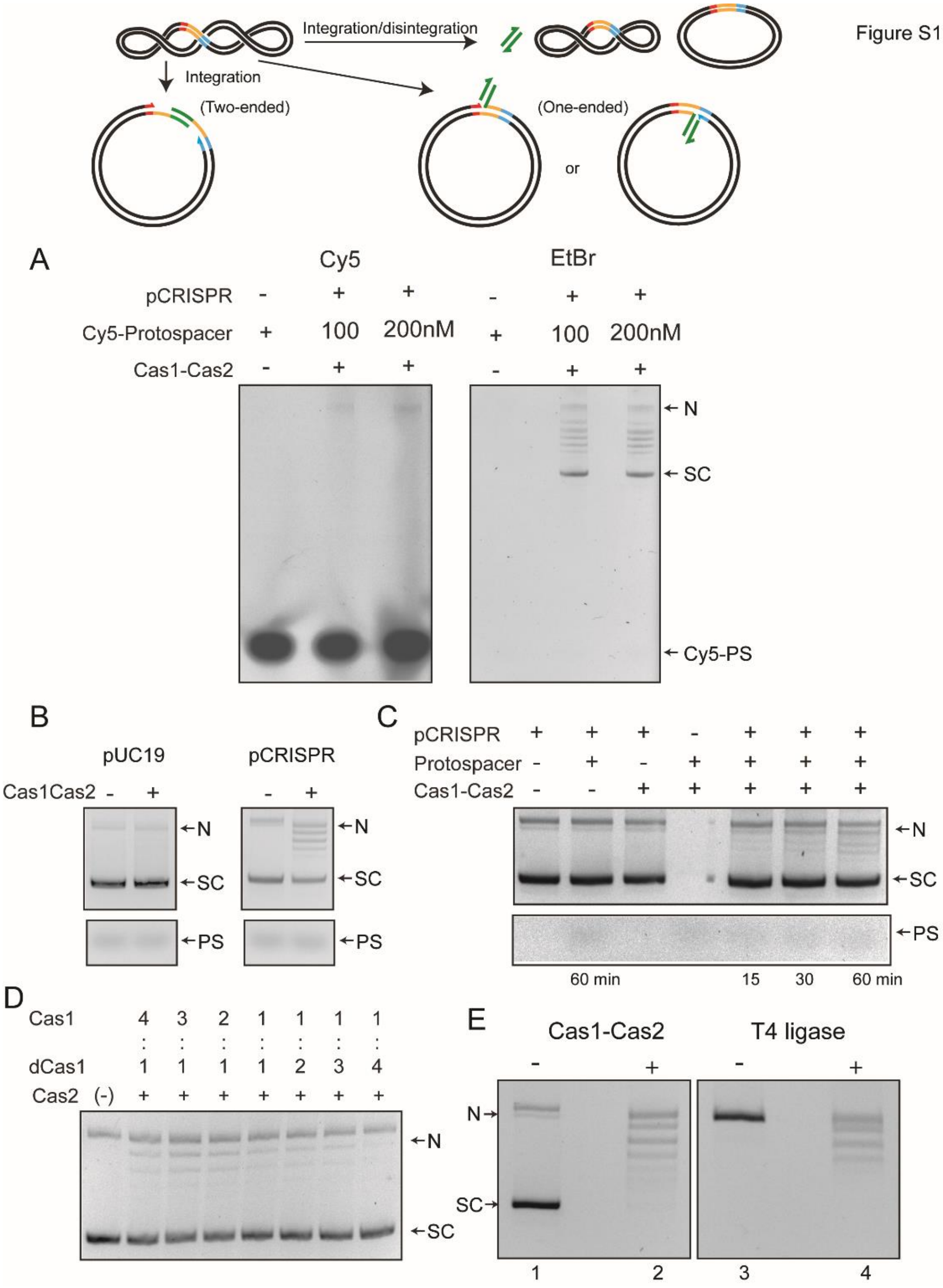

**Figure S1**. **Integration-disintegration by *S. pyogenes* (Spy) Cas1-Cas2 is associated with plasmid relaxation**. **A**. The reactions were performed using the same target site-containing supercoiled plasmid substrate as that used for the assays shown in Figure 1. The proto-spacer was the same as well, except that it contained the Cy5 fluorophore at the 5'-end on both strands. The agarose gel in which the reactions were fractionated was analyzed by fluorography (left panel), and was subsequently stained with ethidium bromide to visualize all DNA bands. Note that there was no fluorescence associated with the supercoiled plasmid or the relaxed plasmid bands. Faint fluorescence was detected in a band that migrates at the position of the nicked plasmid. **B**. Reactions utilized the same plasmid as in **A** (pCRISPR) or one lacking the integration target site (pUC19). **C**. Individual components were omitted from the reactions with the pCRISPR plasmid, as indicated. **D**. These reactions contained Cas1 and dCas1 (catalytically inactive due to the H205A mutation) in the indicated molar ratios. The molar ratio of (Cas1 + dCas1) to Cas2 was maintained at 2:1 in all reactions. **E**. Integration-disintegration reactions were performed with Cas1-Cas2 (left panel). The nicked plasmid was ligated in the absence of Cas1-Cas2 using T4 DNA ligase (right panel). The gel was run in the absence of an intercalator and was then stained with ethidium bromide. SC = supercoiled plasmid; N = nicked plasmid; PS = proto-spacer.

Nearly all of the proto-spacer integration events in the target plasmid appear to be one-ended (transfer of only one strand), with integration being rapidly reversed by disintegration to give covalently closed plasmid molecules. Integration-disintegration requires the target site, the proto-spacer and the functional Cas1-Cas2 complex. The strand transfer reaction is performed by the Cas1 active site. The nicked strand resulting from proto-spacer integration is not impeded from rotation before nick closure. As a result, a relaxed distribution of plasmid circles is formed by disintegration.

Figure S2

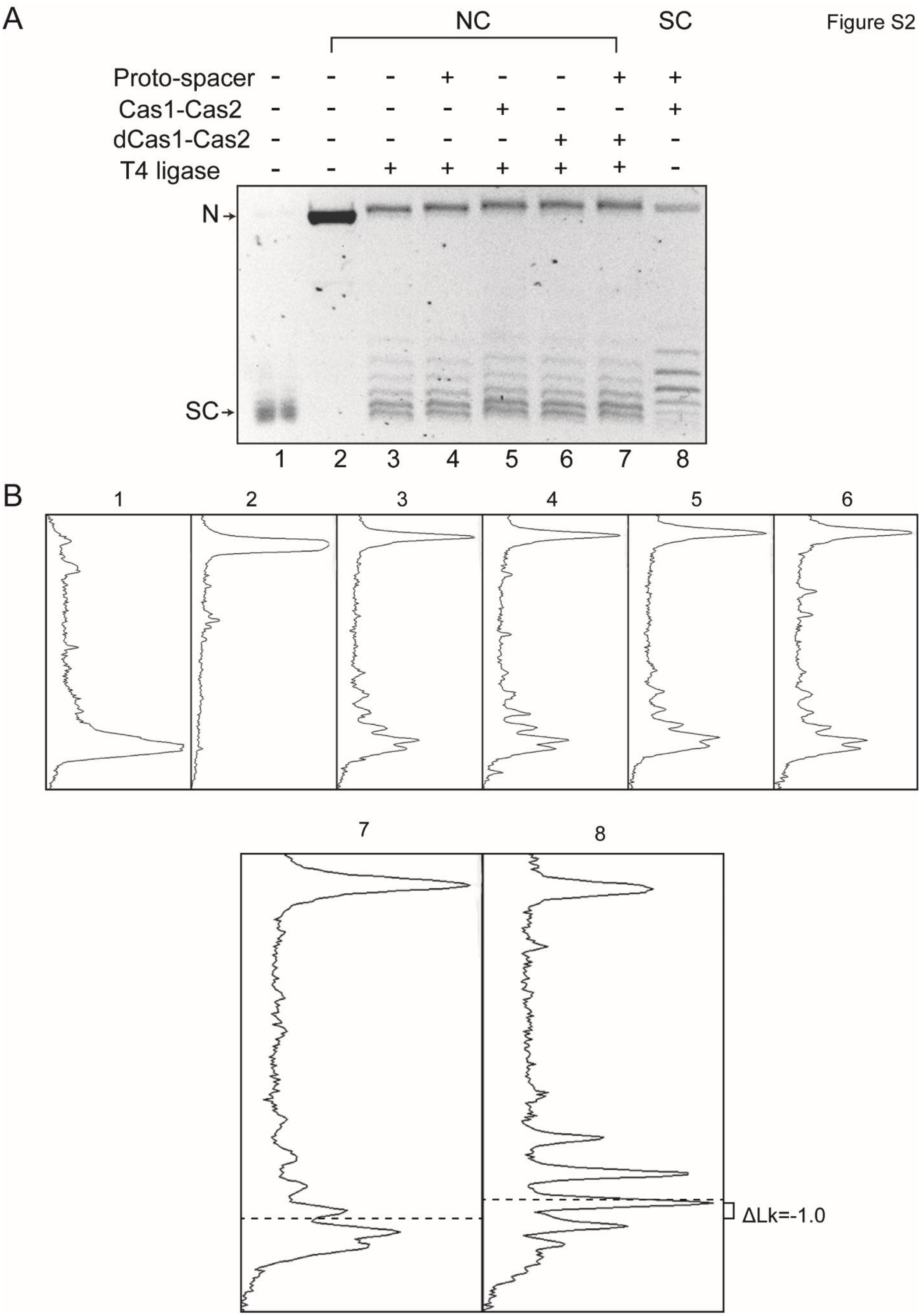

A
NC
SC
Proto-spacer - - - + - - + +
Cas1-Cas2 - - - - + - - +
dCas1-Cas2 - - - - - + + -
T4 ligase - - + + + + + -
N
SC
1 2 3 4 5 6 7 8
B
1
2
3
4
5
6
7
8
ΔLk=-1.0

**Figure S2**. **Cas1-Cas2 unwinds DNA within the target site during proto-spacer integration-disintegration**. **A**. The image of the gel panel shown in Figure 1F (electrophoresis done in the presence of 0.4 μg/ml chloroquine) is reproduced here. **B**. The densitometric tracings of individual DNA bands in each lane are shown in the panels below. The tracings for lane 7 and 8 are enlarged, and the centers of their distributions are marked by the dashed lines.

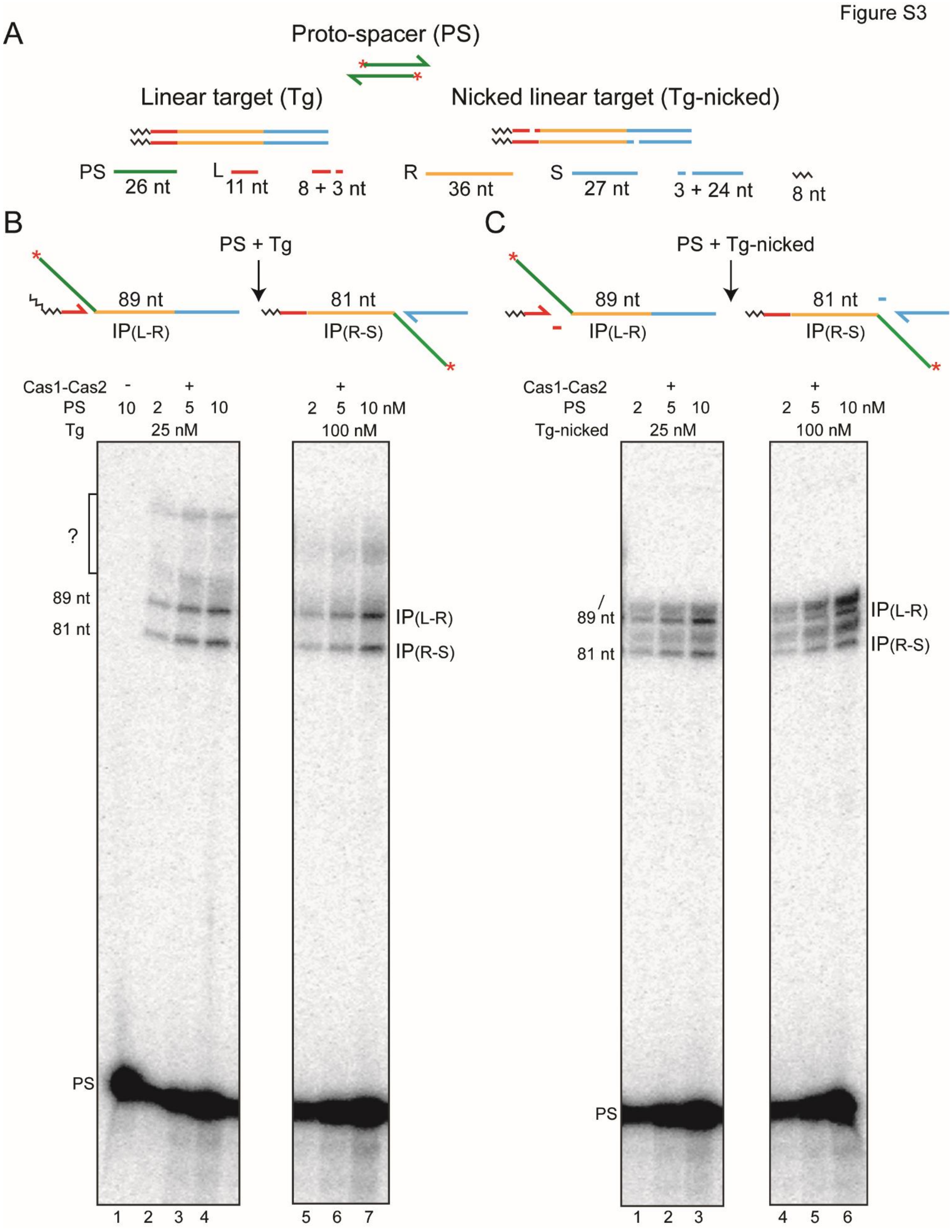
Figure S3
A
Proto-spacer (PS)
Linear target (Tg)
Nicked linear target (Tg-nicked)
PS 26 nt
L 11 nt
8 + 3 nt
R 36 nt
S 27 nt
3 + 24 nt
8 nt
B
PS + Tg
89 nt
IP(L-R)
81 nt
IP(R-S)
Cas1-Cas2 - +
PS 10 2 5 10
Tg 25 nM
+
2 5 10 nM
100 nM
?
89 nt
81 nt
IP(L-R)
IP(R-S)
PS
1 2 3 4
5 6 7
C
PS + Tg-nicked
89 nt
IP(L-R)
81 nt
IP(R-S)
Cas1-Cas2 +
PS 2 5 10
Tg-nicked 25 nM
+
2 5 10 nM
100 nM
89 nt
81 nt
IP(L-R)
IP(R-S)
PS
1 2 3
4 5 6

1

**Figure S3**. **Proto-spacer integration by Cas1-Cas2 in linear target sites is primarily on-target**. An uncropped view of the gel panels corresponding to Figure 2B,C is shown here. The unexpected (and uncharacterized) faint bands above the 89 nt authentic integration product are marked by '?'. Their mobilities suggest off-target integrations within the leader on the top strand or within the spacer on the bottom strand. Prior *in vitro* studies showed that the Spy Cas1-Cas2 complex is more specific in target site selection than *E. coli* Cas1-Cas2 complex (19,33). IP = Integration product; PS = Proto-spacer.

Figure S4

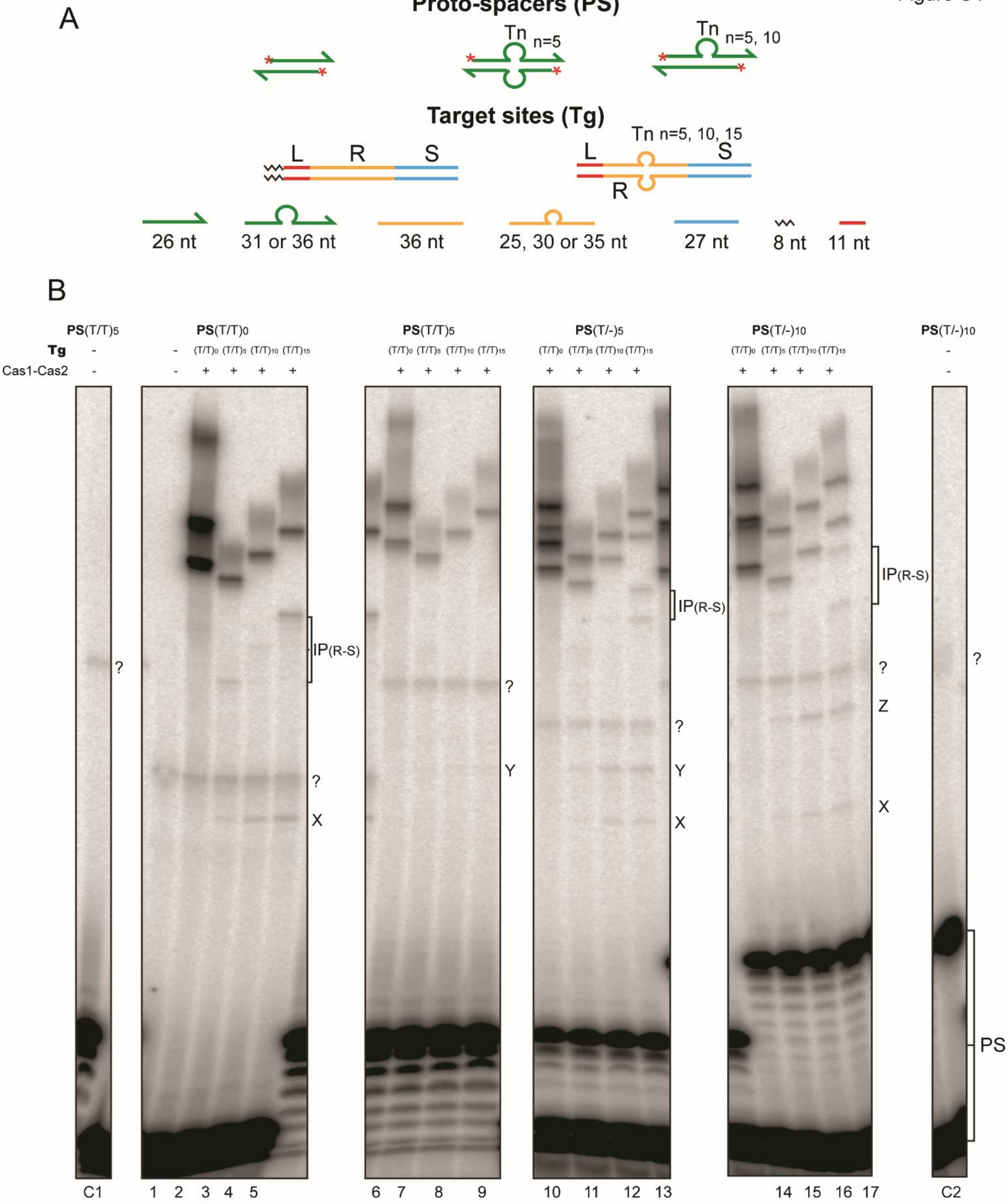

A
Proto-spacers (PS)
Tn n=5
Tn n=5, 10
Target sites (Tg)
L R S
Tn n=5, 10, 15
L S R
26 nt
31 or 36 nt
36 nt
25, 30 or 35 nt
27 nt
8 nt
11 nt
B
PS(T/T)5
PS(T/T)0
PS(T/T)5
PS(T/-)5
PS(T/-)10
PS(T/-)10
Tg
Cas1-Cas2
(T/T)0 (T/T)5 (T/T)10 (T/T)15
IP(R-S)
?
X
Y
Z
PS
C1 1 2 3 4 5 6 7 8 9 10 11 12 13 14 15 16 17 C2

**Figure S4**. **Cas1-Cas2 integrates proto-spacers predominantly at the L-R junction in modified target sites**. **A**. The schematic diagrams of the 5'-end labeled proto-spacers and target sites are arranged as in Figure 3A. **B**. A full view of the source gel for the data for Figure 3B is shown. Phosphorimaging time was extended relative to that in Figure 3B to reveal weak reaction products at the R-S junction, IP(R-S), in the modified targets. For the unmodified proto-spacer $(T/T)_0$, IP(R-S) was now readily detectable in the $(T/T)_{15}$ target (72 nt; lane 5). Even weaker R-S reaction was seen in the $(T/T)_{10}$ target (67 nt; lane 4) and the $(T/T)_5$ target (62 nt; lane 3). Bands for IP(R-S) in the $(T/T)_{15}$ target were also visible for the $(T/-)_5$ proto-spacer (77 nt and 72 nt; lane 13) and the $(T/-)_{10}$ proto-spacer (82 nt and 72 nt; lane 17). For the $(T/-)_5$ proto-spacer and the $(T/T)_5$ or $(T/T)_{10}$ target combinations, IP(R-S) was barely above background (67 nt, 62 nt; lane 11 and 72 nt, 67 nt; lane 12). In all reactions with the modified target sites, the L-R junction far outmatched the R-S junction for insertion efficiency. PS = Proto-spacer.

The bands labeled as X,Y and Z are explained in Text S1 (see below). Unexpected co-migrating sets of bands from reactions of individual proto-spacers with the modified and unmodified target sites, as well as bands that were independent of Cas1-Cas2, are marked with a '?' symbol. These were potential contaminants from the oligonucleotides used for proto-spacer assembly. The incubation mixtures run in lanes C1, 1 and C2 contained the labeled proto-spacer (PS$(T/T)_5$ in C1; PS$(T/T)_0$ in 1; and PS$(T/T)_{10}$ in C2) together with the unmodified target $(T/T)_0$ but no Cas1-Cas2. The C1 and C2 control lanes were not included in the gel panels of Figure 3B. Lanes 1-17 here correspond to lanes 1-17 of Figure 3B.

**Text S1**. The bands X, Y and Z (Figure S4) were formed only in reactions containing the modified $(T/T)_n$ targets, and were dependent on Cas1-Cas2. Their relative spacing (depending on the sizes of the proto-spacer strands) suggest a unique site of insertion common to all the targets. For example, the smallest band X was formed from the native $(T/T)_o$ proto-spacer (with 26 nt on either strand) in all three of the modified targets, $(T/T)_5$, $(T/T)_{10}$ and $(T/T)_{15}$ (Figure S4; lanes 3-5). The mobility of the fainter band Y formed from the $(T/T)_5$ proto-spacer (with 31 nt strands) was consistent with a 5 nt increase in length compared to X (Figure S4; lanes 7-9). With the $(T/-)_5$ proto-spacer (strand lengths of 31 nt and 26 nt), both X and Y were formed (FigureS4; lanes 11-13). And with the $(T/-)_{10}$ proto-spacer (strand lengths of 36 nt and 26 nt), X and Z were formed, the migration of Z suggesting it to be longer than X and Y by 10 nt and 5 nt, respectively (Figure S4; lanes 15-17). Taken together, X, Y and Z appear to be formed by insertions of the 26 nt, 31 nt and 36 nt proto-spacer strands, respectively, at the leader-proximal junction between the $T_n$-insertion and the repeat sequence on the bottom strand. The sizes of the X, Y and Z bands would then be 47 nt, 52 nt and 57 nt, respectively, and match their observed gel mobilities. The double strand-single strand junction may mimic the DNA conformation organized by Cas1-Cas2 at the L-R or R-S junction to promote proto-spacer insertion during the native reaction.

Figure S5

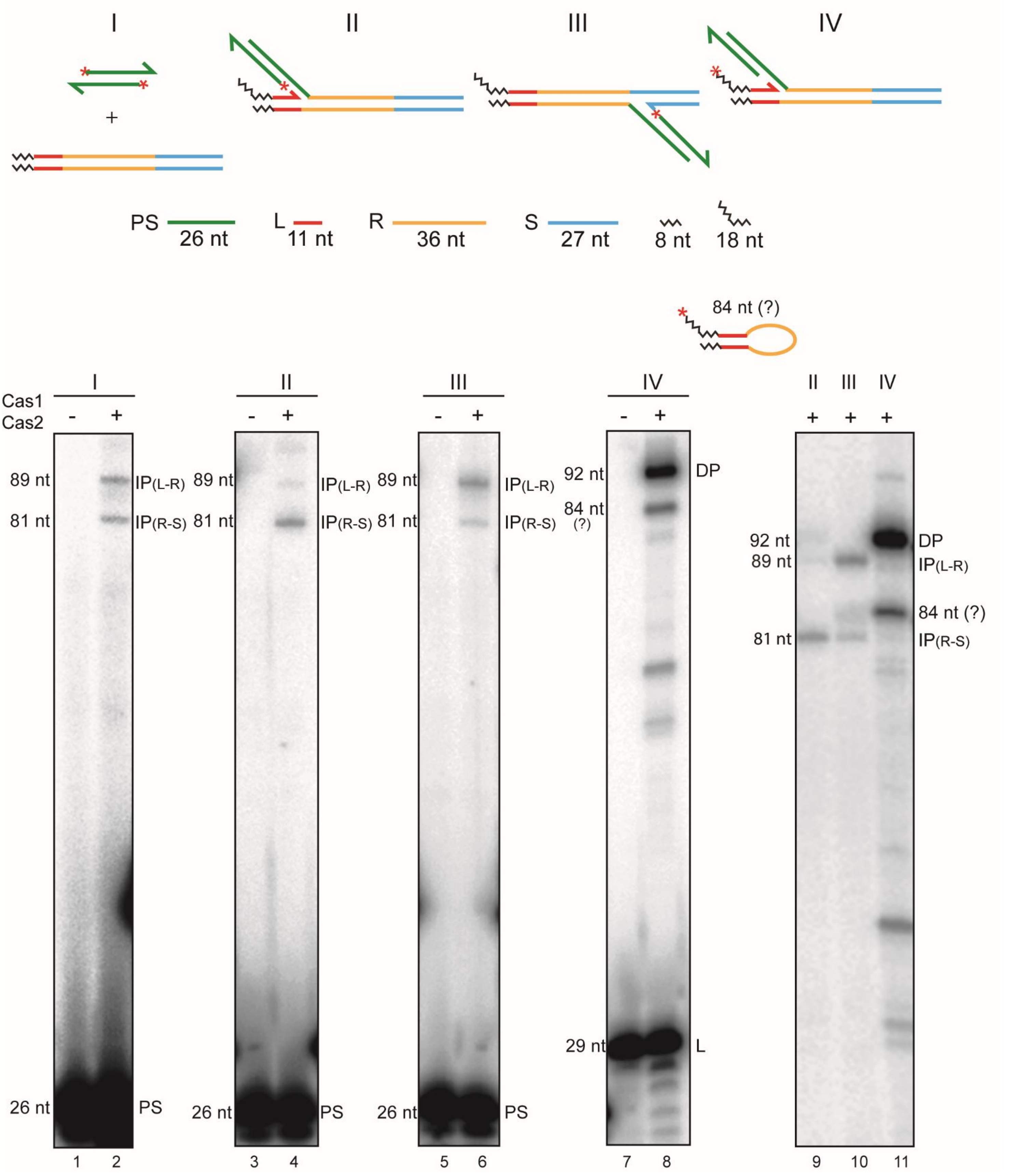

I
II
III
IV
+
PS
26 nt
L
11 nt
R
36 nt
S
27 nt
8 nt
18 nt
84 nt (?)
Cas1
Cas2
-
+
89 nt
81 nt
26 nt
IP(L-R)
IP(R-S)
PS
92 nt
84 nt
(?)
29 nt
DP
L
84 nt (?)
1
2
3
4
5
6
7
8
9
10
11

**Figure S5**. **Cas1-Cas2 mediates integration and disintegration of a semi-integrated proto-spacer**. The substrates for the standard proto-spacer integration reaction and those for the second strand transfer or the disintegration reaction from a semi-integrated proto-spacer are schematically illustrated at the top. The 5'-end label ($^{32}P$) placed on the proto-spacer strands (I), the unintegrated strand of a semi-integrated proto-spacer (II and III), or on the leader strand (IV) is signified by the asterisks. The integration products from a proto-spacer in a linear target site at the L-R junction (lane 2; 89 nt) and the R-S junction (lane 2; 81 nt) provide the reference frame for interpreting the product bands from the 'semi-integrated' substrates. The primary products from the L-R and R-S semi-integrated substrates were integrations of the second proto-spacer strand at the R-S junction (lane 4; 81 nt) and the L-R junction (lane 6; 89 nt), respectively. The weaker 89 nt (in lane 4) and 81 nt (in lane 6) bands are consistent with the reintegration of a protospacer (following its release by disintegration) at the L-R and R-S junctions, respectively. These junctions would be re-formed during disintegration mediated by the 3'-hydroxyls of the leader and spacer strands, respectively. When the $^{32}P$-label was present at the 5'-end of the leader strand, the prominent product was the resealed L-R junction (via disintegration) (92 nt; lane 8). The migration of the weaker band below in lane 8 (see also Figure 5A; lanes 2'-6') is consistent with a size of 84 nt (Text S2 below). Its mobility difference from the 81 nt band (IP(R-S)) was more conspicuous in a longer run gel in which the labeled proto-spacer and leader strands were run off the gel bottom (lanes 9-11). P = Integration product; DP = disintegration product; PS = proto-spacer; L = Leader.

**Text S2**. An 84 nt product is expected if the leader strand were to be transferred to the R-S junction on the opposite strand in an aberrant integration-like reaction. This reaction is formally similar to 'hairpinning' carried out by the Rag1-Rag2 recombinase/transposase and the transposases of the cut-and-paste transposons Tn5 and Tn10 (47-49). Bringing the leader 3'-hydroxyl in proximity to the R-S junction would require considerable distortion of the repeat DNA or its reconfiguration into a cruciform-like structure (19)

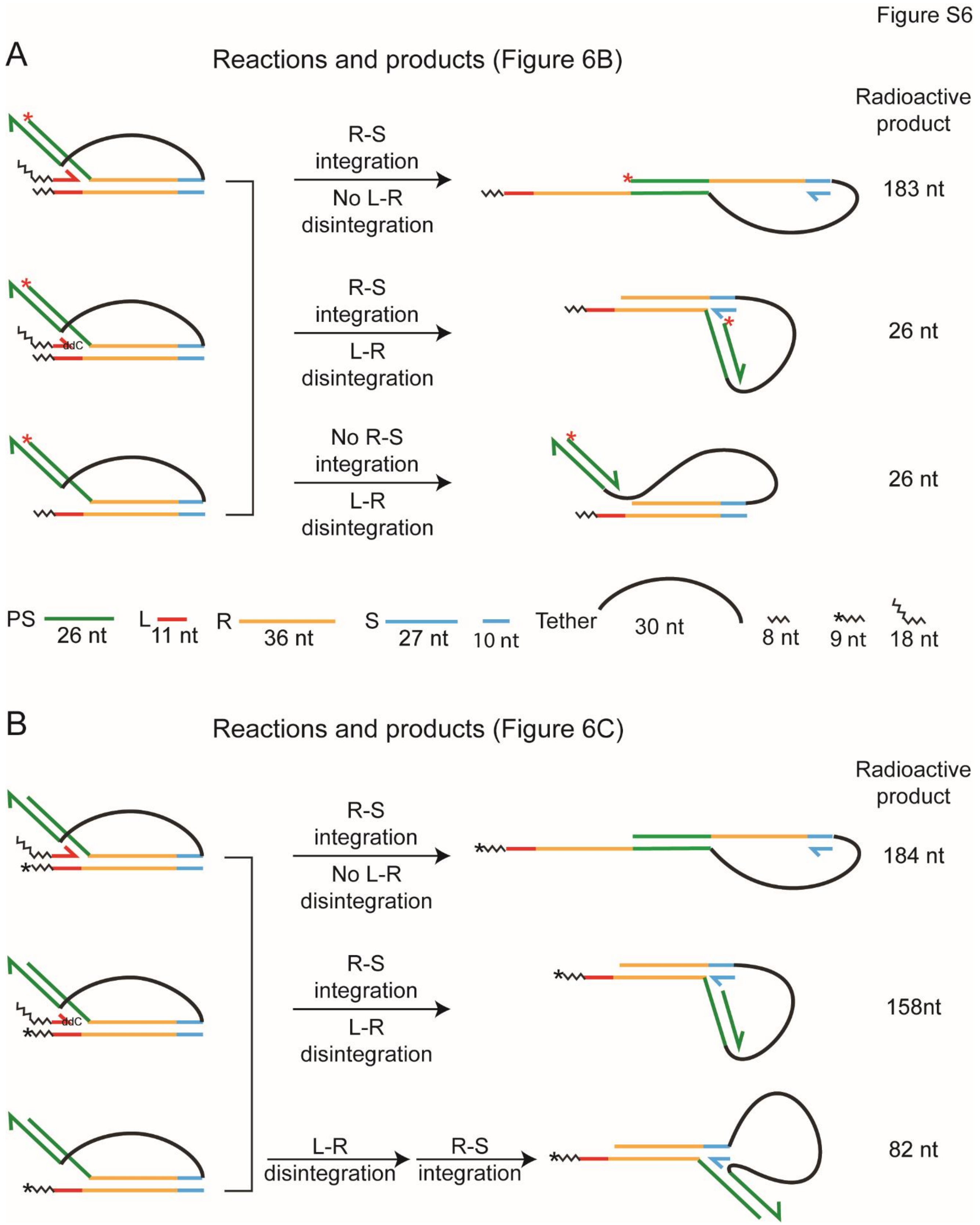

Figure S6
A
Reactions and products (Figure 6B)
Radioactive product
R-S integration
No L-R disintegration
183 nt
ddC
R-S integration
L-R disintegration
26 nt
No R-S integration
L-R disintegration
26 nt
PS
26 nt
L
11 nt
R
36 nt
S
27 nt
10 nt
Tether
30 nt
8 nt
9 nt
18 nt
B
Reactions and products (Figure 6C)
Radioactive product
R-S integration
No L-R disintegration
184 nt
ddC
R-S integration
L-R disintegration
158nt
L-R disintegration
R-S integration
82 nt

1

**Figure S6**. **Distinct products formed by Cas1-Cas2 action on trombone substrates**. The observed reaction products (Figure 6B and C) are schematically illustrated, and the steps involved in their formation are indicated. The unlabeled leader strand is omitted from the figures of the products. The $^{32}$P-label (5'-end; red asterisk) in the substrates in **A** is positioned to identify proto-spacer integration at the R-S junction without its disintegration from the L-R junction. In addition, disintegration of the proto-spacer from the L-R junction (which may or may not be coupled to integration at the R-S junction) is also reported by these substrates. The labeling position in **B** (one $^{32}$P labeled nucleotide at the 3'-end; asterisk) distinguishes integration events at R-S occurring with or without disintegration at L-R. In principle, disintegration and integration may occur in a concerted or sequential manner. The sequential reaction is reported by the 82 nt labeled product, signifying the initial release of the proto-spacer strand inserted at L-R and its subsequent insertion at R-S.

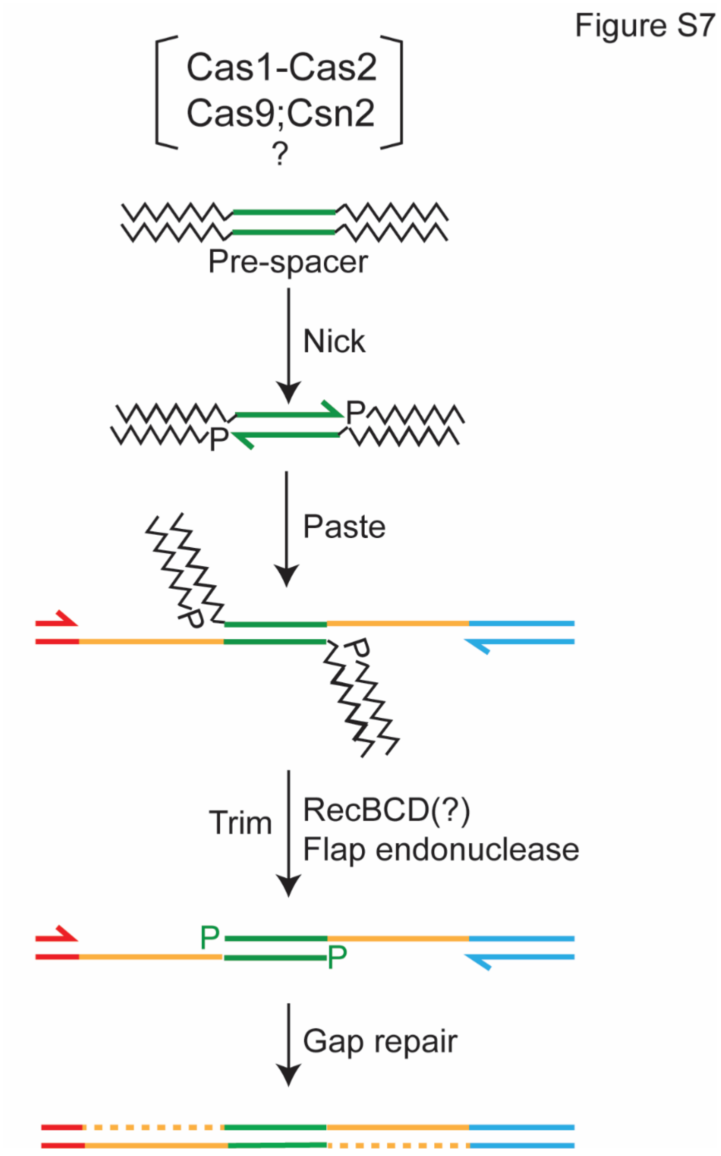


**Figure S7. Spacer acquisition by a mechanism that combines aspects of replicative and cut-and-paste transposition mechanisms**. An integration event may be initiated using the single stranded ends generated by Cas1 activity without the proto-spacer being excised from the pre-spacer DNA (as in replicative transposition; see Figure 7). The flanking DNA is then trimmed by a nuclease such as RecBCD, perhaps with assistance from a flap endonuclease, to generate a structure that resembles the intermediate of cut-and-paste transposition. In effect, the mechanism is ‘nick-paste-and-trim’. The final integration product is formed by gap-filling replication and ligation. The process would be analogous to the removal of host DNA attached to the ends of the infecting phage Mu genome after it has been partially integrated into the *E. coli* chromosome (46).

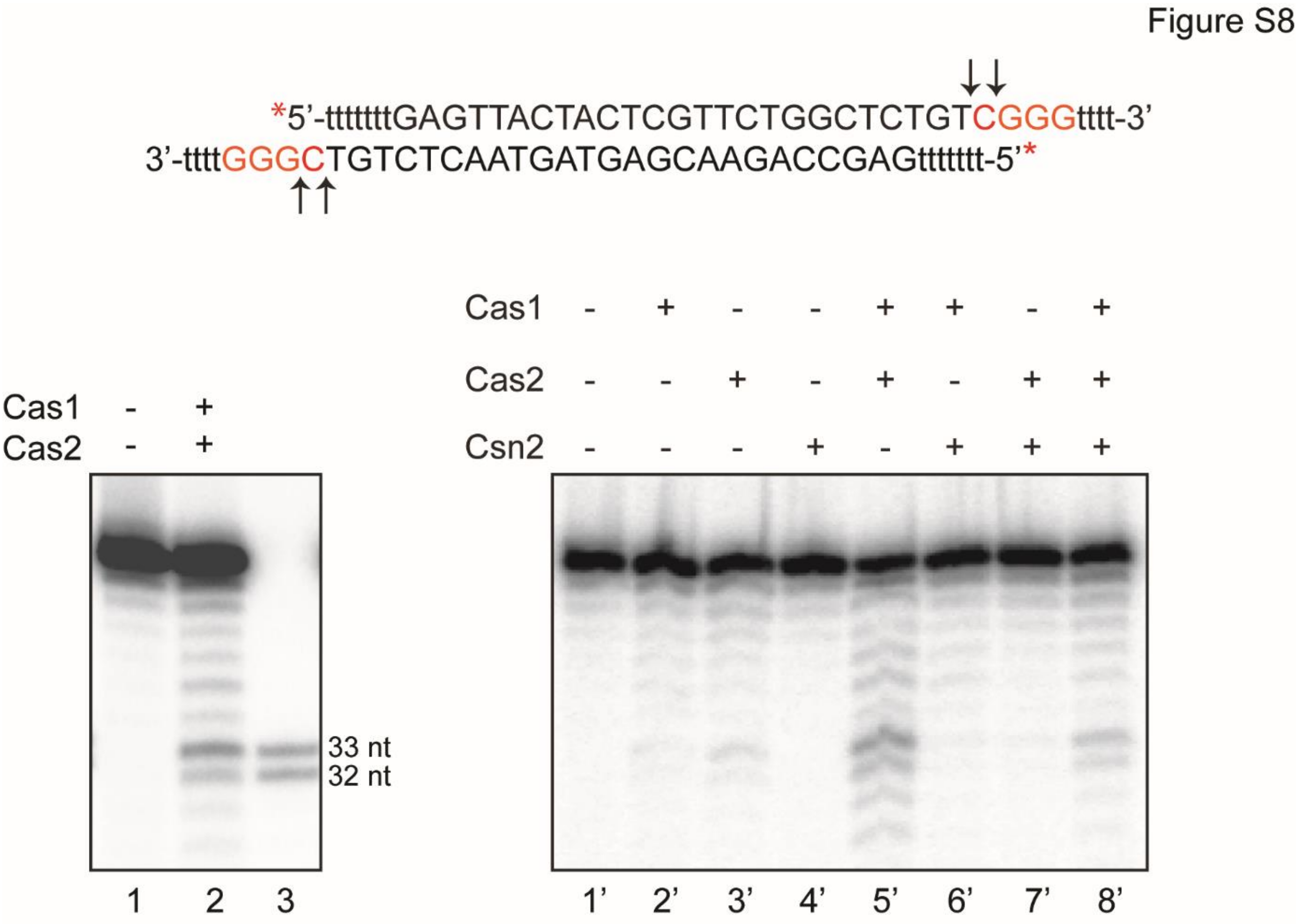


**Figure S8**. **PAM-specific strand cleavage by Cas1-Cas2**. The model pre-spacer DNA substrate contained a 22 bp core flanked on each side by a 7 nt non-complementary double stranded segment, and a 4 nt single stranded 3' extension on either strand. The two flanking segments were identical in sequence. Each non-complementary region contained two overlapping PAM sequences (5'C<u>GG</u>3' and 5'<u>GG</u>G3'; shown in red letters). The cleavage reactions were carried out using 5'-end labeled substrate ($^{32}$P on both strands) and the indicated proteins in the same buffer as that employed for integration assays. Two bands (32 nt and 33 nt) that were prominent over the background cleavage bands in the reactions containing Cas1-Cas2 (lanes 2 and 5') and Cas1-Cas2-Csn2 (lane 8') resulted from cleavage immediately next to the PAM sequences. The cleavage positions are indicated in the substrate sequence by short vertical arrows (5'↓C↓GGG3').

**Text S3**. If PAM-specific cleavage by Cas1-Cas2 is coordinated with transfer of this cleaved end to the L-R junction, the inserted spacer will be in the correct orientation to generate the functional crRNA. This coupled strand cleavage and transfer is easier to imagine in a high-order Cas protein complex capable of both pre-spacer processing and proto-spacer insertion than in a Cas1-Cas2 complex associated with an already processed proto-spacer. *In vitro*, either of the two proto-spacer strands is equally capable of integration at L-R. The problem may be potentially circumvented if processing occurs only when two appropriately spaced PAM sequences are recognized on the two strands of the pre-spacer. In this case, insertion in either orientation would give immunity. For a PAM of the type 5'NGG3', a phage or plasmid genome large enough to be potentially harmful to the host bacterium is likely to contain at least one, and probably several, functional proto-spacers.

| Oligomer | Sequence (5' to 3') | Relevant figures |
|---|---|---|
| Proto-spacer | GAGTTACTACTCGTTCTGGCTCTGTC<br><br>GAGCCAGAACGAGTAGTAACTCTGTC | Figures 1, 2, 3, 4, S1, S3, S4 and S5 |
| Target site | CGATCGATTAGTCTACGAGGTTTTAGAGCTATGCTGTTTTGAATG GTCCCAAAACTGCGCTGGTTGATTTACATGTCTCTCT<br><br>AGAGAGACATGTAAATCAACCAGCGCAGTTTTGGGACCATTCAAA ACAGCATAGCTCTAAAACCTCGTAGACTAATCGATCG | Figures 2, 3, 4, S3, S4 and S5 |
| Target sites (nicked) | CGATCGATTAGTCTAC<br><br>GAGGTTTTAGAGCTATGCTGTTTTGAATGGTCCCAAAACTGCGCT GGTTGATTTACATGTCTCTCT<br><br>AGAGAGACATGTAAATCAACCAGC<br><br>GCAGTTTTGGGACCATTCAAAACAGCATAGCTCTAAAACCTCGTA GACTAATCGATCG | Figures 2 and S3 |
| Proto-spacers with (T)n insertions | GAGTTACTACTtttttCGTTCTGGCTCTGTC<br><br>GAGCCAGAACGtttttAGTAGTAACTCTGTC<br><br>GAGTTACTACTttttttttttCGTTCTGGCTCTGTC<br><br>GAGTTACTACTttttttttttCGTTCTGGCTCTGTC | Figures 3 and S4 |
| Target sites with (T)n insertions | TAGTCTACGAGGTTTTAGAGCtttttGTCCCAAAACTGCGCTGGTTGA TTTACATGTCTCTCT<br><br>AGAGAGACATGTAAATCAACCAGCGCAGTTTTGGGACtttttGCTCTA AAACCTCGTAGACTA<br><br>TAGTCTACGAGGTTTTAGAGCttttttttttGTCCCAAAACTGCGCTGGTT GATTTACATGTCTCTCT<br><br>AGAGAGACATGTAAATCAACCAGCGCAGTTTTGGGACttttttttttGCT CTAAAACCTCGTAGACTA<br><br>TAGTCTACGAGGTTTTAGAGCtttttttttttttttGTCCCAAAACTGCGCTG GTTGATTTACATGTCTCTCT<br><br>AGAGAGACATGTAAATCAACCAGCGCAGTTTTGGGACtttttttttttttttG CTCTAAAACCTCGTAGACTA | Figures 3 and S4 |

| | | |
|---|---|---|
| Half-target site (L-R). | CGATCGATTAGTCTACGAGGTTTTAGAGCTATGCTGT<br><br>ACAGCATAGCTCTAAAACCTCGTAGACTAATCGATCG | Figure 4 |
| Half-target site (R-S) | TTTGAATGGTCCCAAAACTGCGCTGGTTGATTTACATGTCTCTCT<br><br>AGAGAGACATGTAAATCAACCAGCGCAGTTTTGGGACCATTCAAA | Figure 4 |
| Half-target site (L-R, +4) | CGATCGATTAGTCTACGAGGTTTTAGAGCTATGCTGTTTTG<br><br>CAAAACAGCATAGCTCTAAAACCTCGTAGACTAATCGATCG | Figure 4 |
| Half-target site (R-S, +4) | CTGTTTTGAATGGTCCCAAAACTGCGCTGGTTGATTTACATGTCT CTCT<br><br>AGAGAGACATGTAAATCAACCAGCGCAGTTTTGGGACCATTCAAA ACAG | Figure 4 |
| Half-target site (R-S, +6) | TGCTGTTTTGAATGGTCCCAAAACTGCGCTGGTTGATTTACATGT CTCTCT<br><br>AGAGAGACATGTAAATCAACCAGCGCAGTTTTGGGACCATTCAAA ACAGCA | Figure 4 |
| Semi-integration mimics | GAGTTACTACTCGTTCTGGCTCTGTCGTTTTAGAGCTATGCTGTTT TGAATGGTCCCAAAACTGCGCTGGTTGATTTACATGTCTCTCT<br><br>GAGCCAGAACGAGTAGTAACTCTGTC<br><br>ATCGATCGAT CGATCGATTAGTCTACGAG<br><br>AGAGAGACATGTAAATCAACCAGCGCAGTTTTGGGACCATTCAAA ACAGCATAGCTCTAAAACCTCGTAGACTAATCGATCG<br><br>ATCGATCGATCGATCGATTAGTCTACGAC/3ddC<br><br>GAGTTACTACTCGTTCTGGCTCTGTCGTTTTAGAGCTATGCTGTTT TGAATGGTCCCAAAACTGCGCTGGTT<br><br>AACCAGCGCAGTTTTGGGACCATTCAAAACAGCATAGCTCTAAAA CCTCGTAGACTAATCGATCG<br><br>GAGTTACTACTCGTTCTGGCTCTGTCGTTTTAGAGCTATGCTGTTT TGAATGGTCCCAAAACTGCGCTGGTTTTTTTCTCTCTTTTTTGAGC CAGAACGAGTAGTAACTCTGTC | Figures 5, 6 and S5<br><br>Figures 5 and 6<br><br>Figure 6 |

| | | |
|---|---|---|
| Semi-integration mimics (continued) | GAGTTACTACTCGTTCTGGCTCTGTCGTTTTGGGACCATTCAAAACAGCATAGCTCTAAAACCTCGTAGACTAATCGATCG<br><br>GAGCCAGAACGAGTAGTAACTCTGTC<br><br>ATCGATCGATAGAGAGACATGTAAATCAACCAGCGCA<br><br>CGATCGATTAGTCTACGAGGTTTTAGAGCTATGCTGTTTTGAATGGTCCCAAAACTGCGCTGGTTGATTTACATGTCTCTCT | Figure S5 |
| Proto-spacer (for Cas1-Cas2 cleavage) | TTTTTTTGAGTTACTACTCGTTCTGGCTCTGTCGGGTTTT<br><br>TTTTTTTGAGCCAGAACGAGTAGTAACTCTGTCGGGTTTT | Figure S8 |

**Table S1. Synthetic oligonucleotides**. The sequences of the oligonucleotides used for this study are listed. The lower case ‘t’s refer to thymine insertions that remain bulged (with no paired or unpaired bases on the opposite strand) or double-looped (unpaired thymine bases on the opposite strand) in the assembled substrates. The ‘dd’ abbreviation refers to ‘dideoxy’. The assays in which the oligos were used are indicated by the corresponding figure numbers.
